# The kinesin-4 family member KIF27 regulates mitotic progression, cytokinesis and genome stability

**DOI:** 10.64898/2026.06.26.734704

**Authors:** Sascha Pust, Simona M. Migliano, Andreas Brech, Stefan G. Stanciu, Harald Stenmark, Kaisa Haglund

**Affiliations:** Department of Molecular Cell Biology, Institute for Cancer Research, Oslo University Hospital, Montebello, 0379 Oslo, Norway; Centre for Cancer Cell Reprogramming, Institute of Clinical Medicine, Faculty of Medicine, University of Oslo, Montebello, 0379 Oslo, Norway; Center for Microscopy-Microanalysis and Information Processing, National University of Science and Technology POLITEHNICA Bucharest, 060042 Bucharest, Romania; Photon-X Spectrum Lab, CAMPUS Research Institute, National University of Science and Technology POLITEHNICA Bucharest, 060042 Bucharest, Romania

**Keywords:** KIF27, kinesin-4, mitosis, cytokinesis, centralspindlin, ESCRT, abscission, midbody, nuclear morphology

## Abstract

Kinesins are microtubule-dependent motors, yet the functions of the kinesin-4 family member KIF27 remain poorly understood. Here, we demonstrate a dynamic and cell-cycle-dependent localization of KIF27, consistent with its functional roles in mitotic progression. Upon mitotic entry, KIF27 relocates to condensed chromosomes. During anaphase, a fraction of KIF27 accumulates at the spindle midzone, and in telophase and late stages of cytokinesis it localizes at the midbody, colocalizing with key cytokinetic regulators at both structures. Recruitment of KIF27 to the midbody depends on KIF23 and CEP55. KIF27 depletion results in profound cell division defects, altered midbody and microtubule morphology, delayed cytokinesis and cytokinesis failure. Beyond cell division, KIF27 depletion directly compromises nuclear morphology, and pan-cancer transcriptomic analyses correlate low KIF27 expression with aneuploidy and poor patient survival in several cancer types. Together, our results identify KIF27 as a novel regulator of mitotic fidelity and genome stability.

## Introduction

Cell division is one of the most essential and tightly controlled processes in biology, involving the precise separation of genetic material and physical partitioning of daughter cells (Glotzer, 2005). Ensuring accuracy during chromosome segregation in mitosis and subsequent splitting of the cytoplasm through cytokinesis is critical for maintaining a stable genome. Disruptions in these steps can result in aneuploidy, multinucleation, and genomic instability, which are well-known features of both cancer and developmental disorders (Lens and Medema, 2019; Tanaka and Hirota, 2009).

During early anaphase, the central spindle is formed, which dictates the position of the cleavage furrow (Glotzer, 2005). The centralspindlin complex, a heterotetramer of the kinesin-6 motor protein KIF23 (MKLP1) and the Rho-GTPase-activating protein RacGAP1 (also known as MgcRacGAP/CYK-4), is a master organizer of the central spindle (Pan et al., 2021; White and Glotzer, 2012). Centralspindlin bundles antiparallel microtubules at the spindle midzone and recruits effectors necessary for cytokinesis and abscission (Hutterer et al., 2009; Lekomtsev et al., 2012). Abscission is the final stage of cytokinesis, during which the intercellular bridge is severed to complete the physical separation of the daughter cells (Andrade and Echard, 2022). This process requires multiple coordinated protein machineries, including the Endosomal Sorting Complex Required for Transport (ESCRT) (Agromayor and Martin-Serrano, 2013; Vietri et al., 2020). The microtubule-severing enzyme Spastin also plays a critical role in dismantling microtubules in the bridge (Lumb et al., 2012). The midbody protein CEP55 recruits the scaffold protein ALIX and ESCRT-I/TSG101, which coordinately recruit ESCRT-III/CHMP4B to the midbody that in turn promotes membrane scission of the thin intercellular bridge (Carlton et al., 2008; Carlton and Martin-Serrano, 2007; Christ et al., 2016; Guizetti et al., 2011; Morita et al., 2007; Zhao et al., 2006). Mitotic kinases, such as Aurora B, ensure proper timing by activating abscission checkpoints (Petsalaki and Zachos, 2016; Steigemann et al., 2009). In addition, ALIX and syndecan-4/syntenin-1, stabilize ESCRT-III for efficient abscission (Addi et al., 2020; Andrade and Echard, 2022).

Kinesin superfamily proteins represent a diverse group of microtubule-dependent motor proteins that play essential roles throughout cell division (Hirokawa et al., 2009; Wollman et al., 2014; Yildiz, 2025). These ATP-driven motor proteins are divided into 14 distinct families, each responsible for specialized roles, such as intracellular transport, organizing the mitotic spindle, moving chromosomes, and facilitating cytokinesis (Lawrence et al., 2004; Miki et al., 2001).

Chromokinesins are kinesin motor proteins that couple microtubule dynamics with chromatin interactions, thereby controlling chromosome dynamics and segregation fidelity, ensuring genomic stability during cell division (Almeida and Maiato, 2018; Wandke et al., 2012; Zhong et al., 2016). Interference with chromokinesin function can lead to errors in chromosome segregation and genomic instability (Mazumdar and Misteli, 2005). The members of the kinesin-4 (KIF4A), kinesin-10 (KIF22/Kid), and kinesin-12 (KIF15/HKLP2) families are chromokinesins and play critical roles in chromosome positioning that drive chromosome congression and alignment on the mitotic spindle (Almeida and Maiato, 2018; Levesque and Compton, 2001; Takahashi et al., 2016; Vanneste et al., 2011; Wang and Adler, 1995). The Kid motor protein is also maintains proper metaphase spindle size to ensure mitotic fidelity (Tokai-Nishizumi et al., 2005).

The kinesin-4 family in vertebrates comprises six distinct members: KIF4A, KIF4B, KIF7, KIF21A, KIF21B, and KIF27 (Hirokawa et al., 2009). The chromokinesin KIF4A regulates chromosome condensation, central spindle organization and cytokinesis. It directly binds chromosomes to modulate their condensation and interacts with PRC1 to control central spindle length (Dong et al., 2018; Gluszek-Kustusz et al., 2023; Takahashi et al., 2016; Tang et al., 2018). While KIF4A has been characterized in detail, the function of other kinesin-4 family members, particularly KIF27, remains largely unclear.

KIF27 and its paralog KIF7 represent the mammalian orthologs of *Drosophila* Costal2 (Cos2), which has been extensively studied in Hedgehog signaling (Endoh-Yamagami et al., 2009; Sisson et al., 1997). While KIF7 controls mammalian Hedgehog signaling (Cheung et al., 2009; Endoh-Yamagami et al., 2009; Liem et al., 2009), a recent study shows that KIF27 does not regulate this pathway but functions as a microtubule scaffold to maintain transition zone integrity and promote motile cilia assembly (Park et al., 2025). Interestingly, biochemical analysis has revealed that KIF27 exhibits unusual kinesin-4 motor motility, displaying slow or even absent transport capabilities along microtubules *in vitro* (Yue et al., 2018). Defects in the mitotic and cytokinetic machineries can impact genome stability in cancer (Lens and Medema, 2019; Levine and Holland, 2018). Chromosomal instability (CIN), the persistent gain or loss of whole chromosomes or chromosome arms, is a hallmark of aggressive tumors and has been linked to dysfunction of kinesin family members and other mitotic regulators (Lens and Medema, 2019).

Here, we report that KIF27 is required for mitotic progression and cytokinesis. By using comprehensive localization studies, functional analyses, and high-resolution microscopy, we demonstrate that KIF27 exhibits a dynamic subcellular distribution throughout the cell cycle and plays a critical role in regulating chromosome dynamics and cytokinetic abscission. Pan-cancer bioinformatic analyses reveal that KIF27 loss is associated with aneuploidy and that high KIF27 expression is prognostically favorable across multiple human cancer types.

## Results

### KIF27 exhibits a dynamic cellular localization during the cell cycle

To investigate the function of KIF27 in cell division, we first examined its cellular localization throughout the cell cycle in HeLa K cells. Immunofluorescence analyses revealed a dynamic and cell cycle–dependent localization of KIF27 (**Fig. 1**). During interphase, KIF27 is predominantly localized within the nucleus (**Fig. 1A**), where it partially co-localizes with RacGAP1 (**Fig. 1B**). In contrast to RacGAP1, KIF27 was also found in nucleoli, as confirmed by colocalization with nucleolin (**Supplementary Fig. 1A**). In the interphase nucleus, KIF27 also partially co-localized with the transcription factor STAT3 (**Supplementary Fig. 1C**). In line with this nuclear localization, the human KIF27 sequence carries several predicted nuclear localization signals (**Supplementary Fig. 1D**, (Kosugi et al., 2009)).

**Figure 1.**
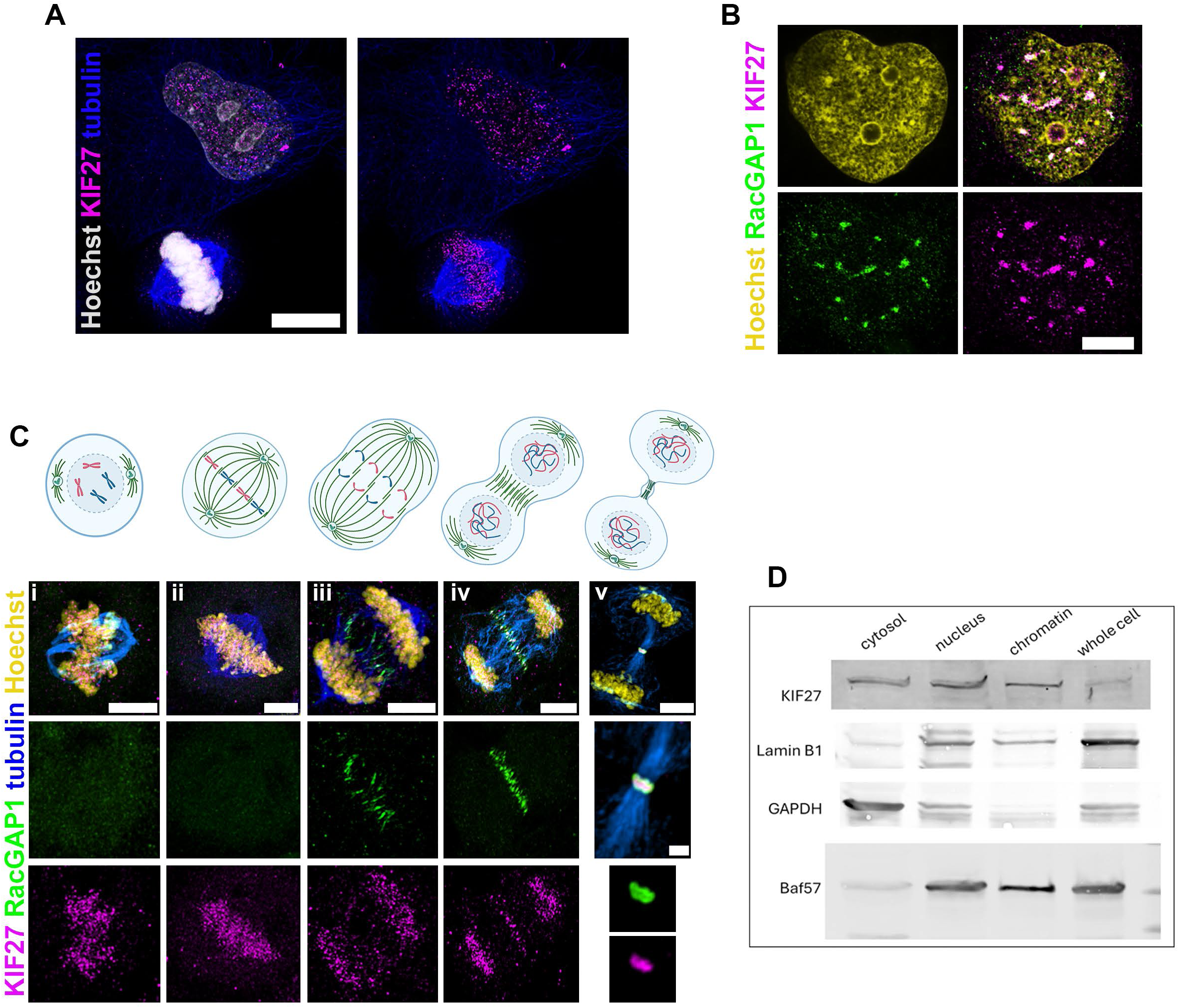
KIF27 exhibits a dynamic cellular localization during the cell cycle. **(A)** Immunofluorescence (IF) images showing the cell cycle-dependent and dynamic localization of KIF27 (magenta). KIF27 is seen in interphase nuclei and associating with chromosomes in mitotic Hela K cells. Microtubules are stained with tubulin (blue), and DNA is stained with Hoechst (white). Scale bar: 10 µm. **(B)** IF images of interphase cells showing KIF27 (magenta) predominantly localized within the nucleus and partially colocalized with RacGAP1 (green). DNA is stained with Hoechst (yellow). Scale bars: 10 µm **(C)** IF images illustrating the dynamic redistribution of KIF27 (magenta) during different mitotic stages and cytokinesis: (i) Prophase/Prometaphase: KIF27 associates with condensed chromosomes. (ii) Metaphase: KIF27 persists on chromosomes at the metaphase plate. (iii, iv) Anaphase: KIF27 persists on segregating chromosomes and begins to accumulate at the spindle midzone (iv). (v) Telophase/Cytokinesis: Strong accumulation of KIF27 at the midbody. RacGAP1 (green), Tubulin (blue), and Hoechst (DNA, yellow) are co-stained. Scale bars: 5 µm (top)and 1 µm (v, inset). **(D)** Western blot of subcellular fractions (cytosol, nucleus, chromatin, whole-cell lysate) showing KIF27 across all fractions with strongest nuclear enrichment. Co-immunoprecipitation of RacGAP1 can be observed in the cytosolic and nuclear fractions. Lamin B1, GAPDH and Baf57 was used to monitor fraction purity.

Upon mitotic entry, KIF27 redistributed from the nucleus to chromosomes and microtubule-associated structures (**Fig. 1C**). During prophase and prometaphase (**Fig. 1C**, i), we observed KIF27 association with condensed chromosomes. In metaphase, chromosomes line up along the metaphase plate, and this was accompanied by an accumulation of KIF27 at the metaphase plate (**Fig. 1C**, ii). During anaphase the sister chromatids separate, and chromosomes are pulled towards the opposite poles of the cell (**Fig. 1C**, iii + iv). The chromosomal KIF27 localization persisted throughout metaphase (**Fig. 1C**, ii) and anaphase (**Fig. 1C**, iii + iv). In late anaphase, a fraction of KIF27 was associated with segregating chromosomes while another KIF27 fraction accumulated in the spindle midzone, where it colocalized with RacGAP1 (**Fig. 1C**, iv). Constriction of the central spindle in telophase leads to the formation of the cytokinetic bridge with a central midbody, consisting of RacGAP1 and KIF23/MKLP1. During telophase, KIF27 accumulated at the midbody, where it colocalized with RacGAP1 (**Fig. 1C**, v).

Combined cell fractionation and Western blot analysis confirmed the presence of KIF27 in both nuclear and chromatin-enriched fractions, supporting its association with nuclear structures. KIF27 was detected in the cytoplasmic, nuclear, and chromatin fractions, with strongest enrichment in the nuclear-associated pools (**Fig. 1D**). The purity of the fractions was monitored with GAPDH, Lamin B1, and Baf57 served as controls for cytosolic, nuclear, and chromatin fractions, respectively (**Fig. 1D**). Like KIF27, KIF4A was detected in all cellular fractions, whereas RacGAP1 was not found in the chromatin fraction (**Supplementary Fig. 1B**).

Collectively, these results demonstrate that KIF27 localizes to both nuclear and cytoplasmic compartments and associates with chromatin, central spindle and midbody structures, suggesting its potential role in chromosome regulation and cytokinesis during cell division.

### KIF27 colocalizes and forms complexes with key cytokinetic regulators of the spindle midzone and midbody

As we observed a highly dynamic redistribution of KIF27 to the spindle midzone and midbody during cell division, we analyzed its relative localization with cytokinetic regulators in greater detail by super resolution microscopy. Importantly, co-staining experiments for KIF27 and the related chromokinesin KIF4A showed KIF27 enrichment around chromosomes during mitosis and then at the midbody during telophase, which partially resembled the distribution of KIF4A (**Fig. 2A**). However, KIF27 and KIF4A showed no significant co-localization on condensed chromosomes, suggesting that the two kinesins occupy distinct chromosomal domains (**Supplementary Fig. 3E**). When we focused on localization of KIF27 in the spindle midzone of anaphase cells, we observed high degrees of colocalization with both centralspindlin components RacGAP1 and KIF23/MKLP1 (**Fig. 2B**) as well as with CEP55 (**Fig. 2C**), which is reported to localize at the spindle midzone prior to midbody localization (Fabbro et al., 2005; Martinez-Garay et al., 2006; Zhao et al., 2006). These key proteins are also involved in the formation and function of the midbody during cytokinesis (Halcrow et al., 2022; Zhao et al., 2006). Co-immunoprecipitation experiments also confirmed an association of KIF27 with RacGAP1 and CEP55 (**Fig. 2D**, left and right panels). However, we could not co-precipitate KIF27 together with KIF23 (**Fig. 2D**, middle panel). STED nanoscopy (Vicidomini et al., 2018) revealed the localization of KIF27 at the interfaces between the microtubules and the midbody in late cytokinesis, with interconnections between the two parts (**Fig. 2E, F** and **Supplementary Movie 1**). Thus, KIF27 emerged as an integral part of the midbody with a characteristic structural organization. Importantly, localization of KIF27 to the midbody was observed consistently across multiple human cell lines (HeLa K, PC3, U2OS and RPE-1), indicating that this recruitment represents a fundamental aspect of its function rather than a cell type-specific phenomenon (**Supplementary Fig. 2A**). Upon observing that KIF27 became highly concentrated at the midbody, we systematically screened for colocalization with key midbody-associated proteins. Optical nanoscopy revealed that KIF27 colocalized with multiple proteins found in the spindle midzone or at the midbody (**Table 1**, and **Supplementary Fig. 2**). Within the midbody, KIF27 clearly colocalized with KIF23 and PRC1 (**Supplementary Fig. 2B** and **2E**), two proteins of the midbody core region (D’Avino and Capalbo, 2016; Hu et al., 2012). Additionally, we performed proximity ligation assays (PLAs), which showed spatial association (≤40 nm) of KIF27 with several midbody-associated proteins, including ALIX, CENP-E and Aurora B, at the midbody (**Table 1** and **Supplementary Fig. 2C**). Interestingly, the KIF27 paralogue KIF7 neither localized to the spindle midzone nor to the midbody, indicating a distinct function for KIF27 in mitosis and cytokinesis (**Supplementary Fig. 2H**). Moreover, no significant colocalization with KIF11, another cytokinetic motor protein at the midbody, was observed (**Supplementary Fig. 2I**).

**Figure 2.**
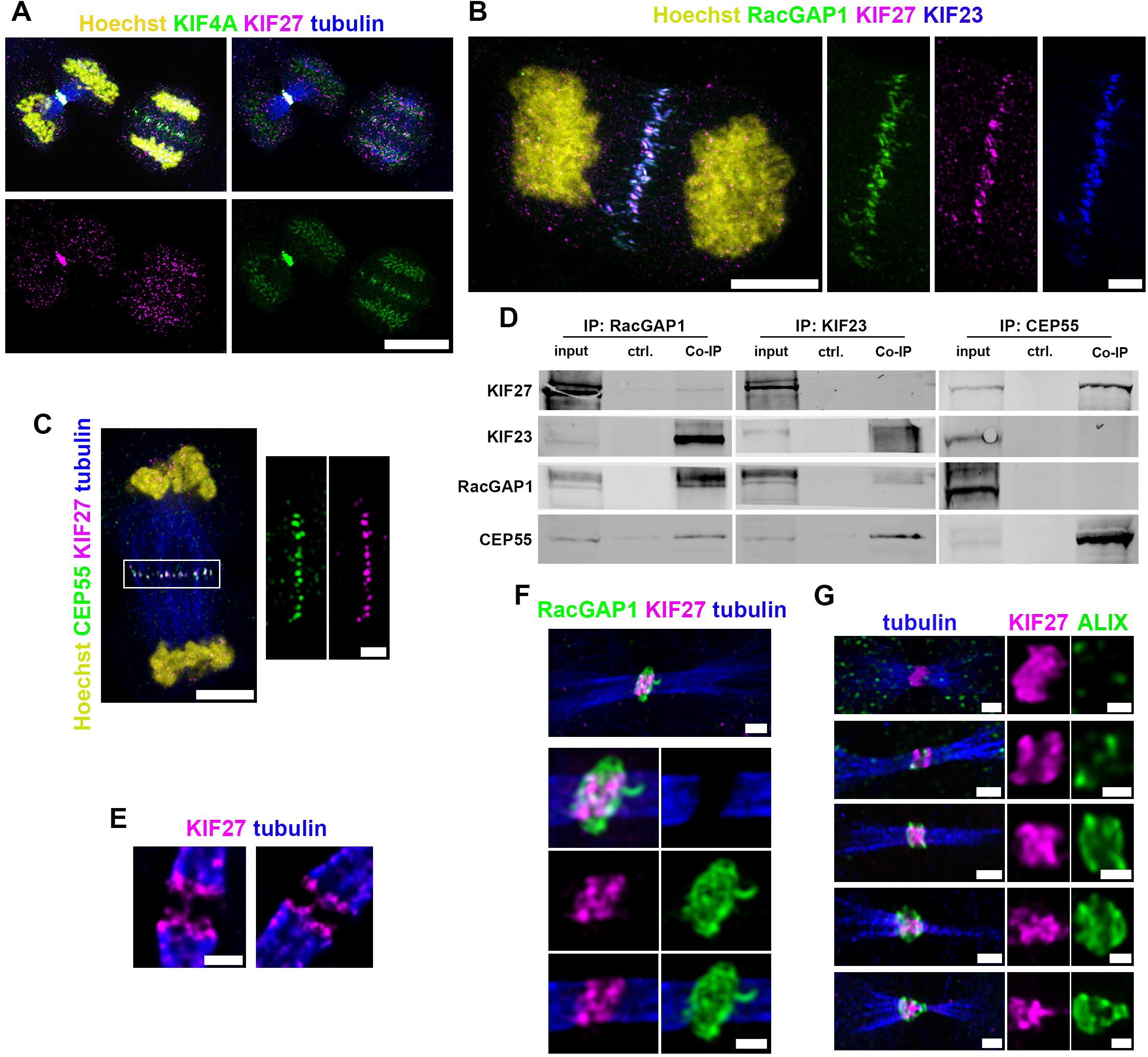
KIF27 colocalizes and forms complexes with key cytokinetic regulators at the spindle midzone and midbody. **(A)** Immunofluorescence (IF) microscopy images showing KIF27 (magenta) localization compared to the related chromokinesin KIF4A (green) during mitosis and cytokinesis. KIF27 enrichment around chromatin during mitosis partially resembles KIF4A distribution. Tubulin (blue) and Hoechst (DNA, yellow) are co-stained. Scale bar: 10µm. **(B + C)** IF images of cells in anaphase indicating significant accumulation of KIF27 (magenta) in the spindle midzone and colocalization with the key regulators of cytokinesis: **(B)** RacGAP1 (green) and KIF23/MKLP1 (blue), and **(C)** CEP55 (green) and tubulin (blue). Hoechst is co-stained in yellow. Scale bars: 5 µm and 2 µm (spindle midzone, insets). **(D)** Co-Immunoprecipitation (Co-IP) experiments showing association of KIF27 with RacGAP1 and CEP55, while KIF23 did not pull down KIF27. Immunoprecipitation (IP) of RacGAP1 led to Co-IP of KIF27, KIF23, and CEP55. Co-IP of RacGAP1 and CEP55 was detected upon IP of KIF23. IP of CEP55 resulted in Co-IP of KIF27. For negative control (ctrl.), uncoupled beads were used. **(E)** STED nanoscopy images showing the subcellular localization of KIF27 (magenta) in late cytokinesis. KIF27 is localized proximal at the interfaces between the microtubules (tubulin, blue) and the midbody, showing interconnections between the exterior parts of both sides. Scale bar: 0.5 µm. **(F)** IF image of KIF27 and RacGAP1 at the midbody of an intercellular bridge. KIF27 (magenta) is localized proximal at the intersection to the microtubule network (blue) and integrated in the centralspindlin complex (RacGAP1, green). Scale bars: 1µm and 0.5 µm (insets). **(G)** Immunofluorescence microscopy images of midbody structures showing KIF27 (magenta), ALIX (green) and tubulin (blue) during distinct progressive phases of cytokinesis, including the development of the intercellular bridge and midbody and the formation of secondary ingressions (from the top to the lower panel). KIF27 accumulates at the midbody prior to ALIX. Both proteins show a dynamic morphological change during bridge and midbody maturation. Scale bars: 1 µm (left panel) and 0.5 µm (insets).

**Table 1.** Summary of proximity studies of KIF27 with different cytokinetic proteins at the spindle midzone or midbody. The data are based on co-localization, co-immunoprecipitation (Co-IP) and proximity ligation assay (PLA) studies (see also Fig. 1D, 2A-C, and suppl. Fig. 2). Proteins are listed alphabetically. +, detected. −, not detected. not tested, not analyzed in this study.

|  | midzone | midbody |
| --- | --- | --- |
| **ALIX** | – | + |
| **Anillin** | not tested | + |
| **Aurora B** | + | + |
| **CENP-E** | + | + |
| **CEP55** | + | + |
| **CHMP4B** | – | + |
| **CITK1** | + | + |
| **KIF4a** | + | + |
| **KIF7** | – | – |
| **KIF11** | – | – |
| **KIF23/MKLP1** | + | + |
| **PRC1** | + | + |
| **RacGAP1** | + | + |

In addition to an early and persistent recruitment KIF27 to the midbody, we observed structural changes of KIF27 morphology during the maturation of the cytokinetic bridge (**Fig. 2G**). Similarly to ALIX, KIF27 transitioned from a disk-like morphology during the early stages of midbody maturation to a cone-like structure during cellular abscission (**Fig. 2G** and **Supplementary Movie 2**). During these stages, KIF27 was detected in the core of the midbody (**Fig. 2G** and **Supplementary Movie 2**).

Taken together, our data show that KIF27 colocalizes with several midzone- and midbody-associated proteins, including KIF23, RacGAP1, PRC1, KIF4A and CEP55 during the progressive stages of cytokinesis. The proximity of KIF27 to these regulators indicates that KIF27 is part of this extensive interaction network and crucial hub within the cytokinetic machinery.

### KIF27 midbody recruitment is dependent on KIF23 and CEP55, but independent of downstream abscission factors

To understand the mechanisms governing KIF27 recruitment during cytokinesis, we performed systematic knockdown experiments targeting key midbody components and analyzed the effects on KIF27 localization. The centralspindlin complex, consisting of RacGAP1 and KIF23/MKLP1, is important for the early recruitment of downstream proteins like CEP55, which in turn mediates midbody recruitment of ALIX and ESCRT-I/TSG101 and ultimately ESCRT-III/CHMP4B (Agromayor and Martin-Serrano, 2013; Andrade and Echard, 2022). After knockdown of either KIF23 or CEP55, the recruitment of KIF27 to the midbody region was clearly decreased (**Fig. 3A-B**). Consequently, midbody recruitment of KIF27 depends on the presence of the centralspindlin protein KIF23 and the early abscission factor recruiter CEP55. In contrast to its dependence on KIF23, depletion of ALIX did not prevent recruitment of KIF27 to the midbody (**Supplementary Fig. 3A-B, Supplementary Movies 3, 4**). We previously described the transport of ALIX to the midbody by a KIF5B-dependent mechanism after the formation of the bridge and midbody (Pust et al., 2023). Consistent with these results, we observed that KIF27 was recruited to the midbody prior to ALIX (**Fig. 2G**).

**Figure 3.**
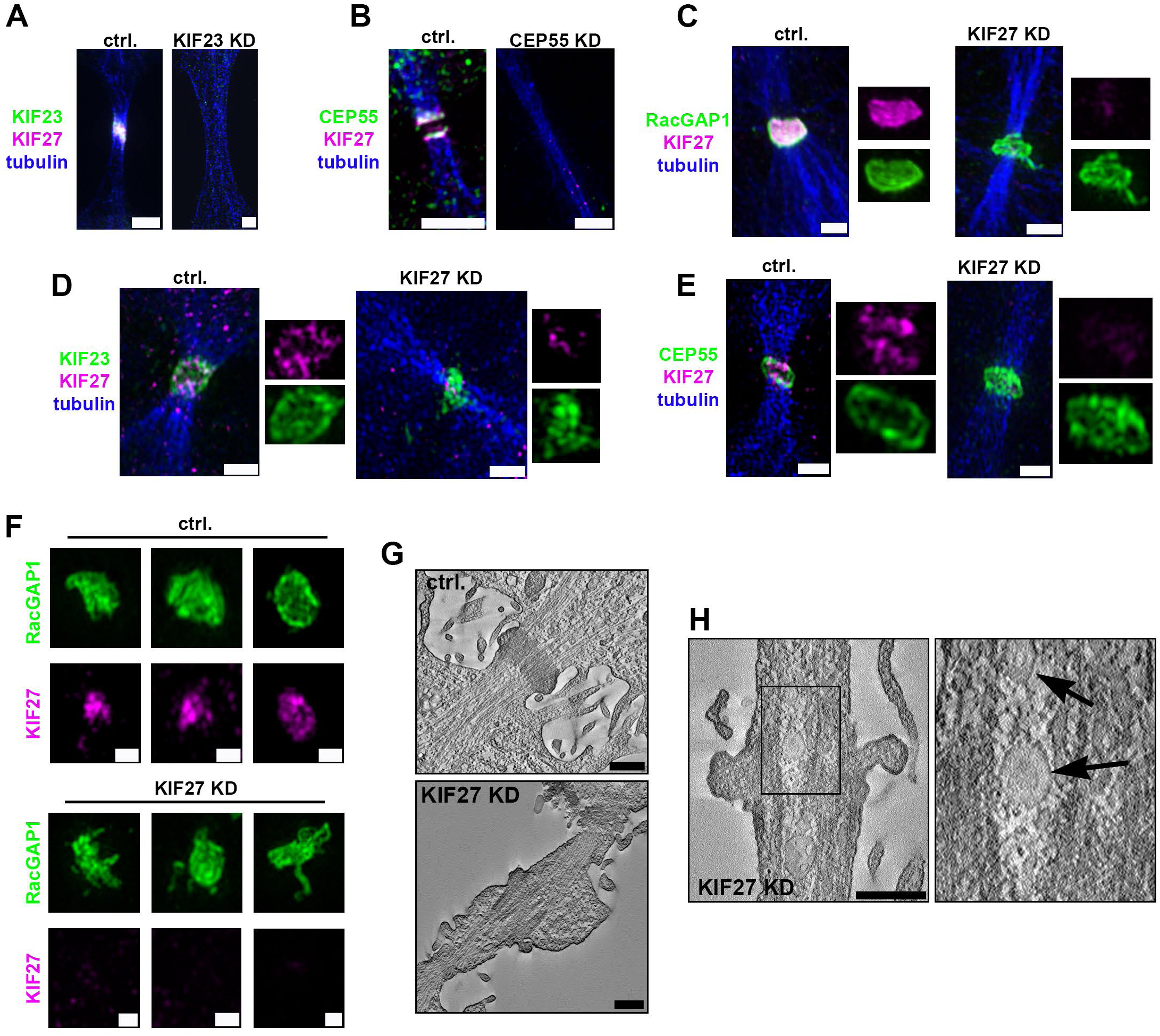
Recruitment of KIF27 to the midbody is dependent on KIF23 and CEP55, and KIF27 is required for structural midbody integrity. **(A +B)** Immunofluorescence (IF) images of KIF27 recruitment to the midbody in control cells (ctrl.) or after siRNA-mediated knockdown (KD) of KIF23 or CEP55. Knockdown of KIF23 **(A)** or CEP55 **(B)** prevents accumulation of KIF27 in the midbody region. KIF27 (magenta), tubulin (blue), and KIF23 (green, **A**) or CEP55 (green, **B**). Scale bars: 2 µm. **(C - E)** IF analysis of the recruitment of key cytokinesis regulators to the midbody upon KIF27 depletion (KD). KIF27 depletion did not inhibit the recruitment of RacGAP1 **(C)**, KIF23 **(D)**, or CEP55 **(E)** to the midbody (green). KIF27 is shown in magenta, tubulin in blue. Scale bars: 1 µm. **(F)** IF analysis (detailed structural analysis) of the centralspindlin complex morphology, stained by RacGAP1 (green), in control (ctrl.) versus KIF27-depleted (KD) cells. Control cells show a condensed ring-like structure, whereas KIF27 depletion resulted in a more diffuse structure with long filamentous protrusions, indicating disruption of the structural midbody integrity. KIF27 is shown in magenta. Scale bars: 500 nm. **(G)** Electron microscopy (EM) tomography demonstrates the morphological defects in the midbody upon KIF27 knockdown (KD). KIF27 KD resulted in a distorted electron-dense midbody pattern in comparison to the uniformly condensed midbody in control cells (ctrl.). **(H)** EM tomography reveals a distorted microtubule network, and the formation of large vesicles in the intercellular bridge (indicated by arrows) upon KIF27 KD, indicating that the structural integrity of the bridge and the midbody were strongly affected. Scale bar: 500 nm.

This finding, combined with our observation that KIF27 is present at the spindle midzone and arrives at the midbody prior to ALIX, strongly suggests that KIF27 functions already in the early stages of midbody maturation, upstream of the ESCRT-dependent abscission machinery.

### KIF27 depletion does not inhibit the recruitment of RacGAP1, KIF23, and CEP55 to the midbody, but interferes with the structural midbody integrity

Next, we analyzed recruitment of RacGAP1, KIF23 and CEP55 to the midbody upon siRNA-mediated depletion of KIF27. Immunofluorescence analysis of the intercellular bridge during cytokinesis revealed that KIF27 depletion neither inhibited the recruitment of RacGAP1, KIF23, nor CEP55 to the midbody (**Fig. 3C-E**). Nevertheless, a detailed structural analysis of the centralspindlin component RacGAP1 revealed disruption of the structural integrity of the midbody (**Fig. 3F**). Control cells showed a condensed RacGAP1-positive midbody structure, whereas KIF27 depletion resulted in irregular midbody structures with long filamentous protrusions (**Fig. 3F**). The morphological defects upon KIF27 knockdown were confirmed by electron microscopy. Control cells showed midbodies with condensed and distinct morphology (**Fig. 3G**, upper image). In contrast, upon KIF27 knockdown, the electron dense midbody appeared distorted (**Fig. 3G**, lower image). Furthermore, STEM (scanning transmission electron microscopy) tomography demonstrated that the structural integrity of the midbody was strongly affected upon KIF27 depletion (**Fig. 3H**). Upon KIF27 knockdown the microtubule network and midbody structure appeared to be distorted, which was sometimes accompanied by the presence of large vesicles in the intercellular bridge (**Fig. 3H** and **Supplementary Movie 5**). These findings indicate that KIF27 has an important role in maintaining the structural integrity of the midbody and the associated microtubule network in the cytokinetic bridge.

### KIF27 depletion causes severe mitotic and cytokinetic defects

To understand the functional importance of KIF27, we analyzed the effects of KIF27 depletion during mitosis and cytokinesis. Live-cell imaging revealed multiple defects upon KIF27 depletion (**Fig. 4** and **Supplementary Fig. 3C**). During 12-14 hours of live-cell imaging we observed a significantly reduced percentage of mitotic cells (**Fig. 4A**; ctrl.: 37.49%, KD: 16.44%) and increased percentage of apoptotic cells upon KIF27 depletion as compared to control cells (**Fig. 4B** and **Supplementary Movie 6**; ctrl.: 1.91%, KD: 7.05%). The percentage of mitotic cells that went through normal and complete cell division was strongly reduced in KIF27-depleted cells (**Fig. 4C** and **Supplementary Movie 7**; ctrl.: 81.06%, KD: 21.84%). KIF27 depletion did not significantly increase the percentage of mitotic cells undergoing irregular cell division (multiple spindle poles, multinucleation; **Fig. 4D** and **Supplementary Movie 8**). However, KIF27 depletion resulted in a large increase of mitotic cells undergoing incomplete mitosis or defective cytokinesis (mitotic arrest, abscission failure; **Fig. 4E** and **Supplementary Movie 9**; ctrl.: 9.79%, KD: 61.18%). During the earlier phases of mitosis, metaphase-to-anaphase transition was prolonged upon KIF27 depletion (**Fig. 4F**; ctrl.: 19.76 min, KD: 32.81 min), which was occasionally accompanied by lagging chromosomes (**Supplementary Movie 10**). Furthermore, in non-synchronized cells we detected an increase in nuclear size of KIF27-depleted cells compared to control cells (**Fig. 4G** and **Supplementary Fig. 3D**), an indication of cell cycle arrest before mitosis. Given the nuclear and nucleolar localization of KIF27 during interphase (**Fig. 1A, B**) and the observed increase in nuclear size upon KIF27 depletion (**Fig. 4G**), we further analyzed nuclear morphology. Nuclear circularity decreased significantly upon KIF27 depletion (**Fig. 4H**), indicating that loss of KIF27 leads to irregular nuclear shape. Taken together, these analyses show that KIF27 is vital to maintain fidelity in mitotic and cytokinetic progression.

**Figure 4.**
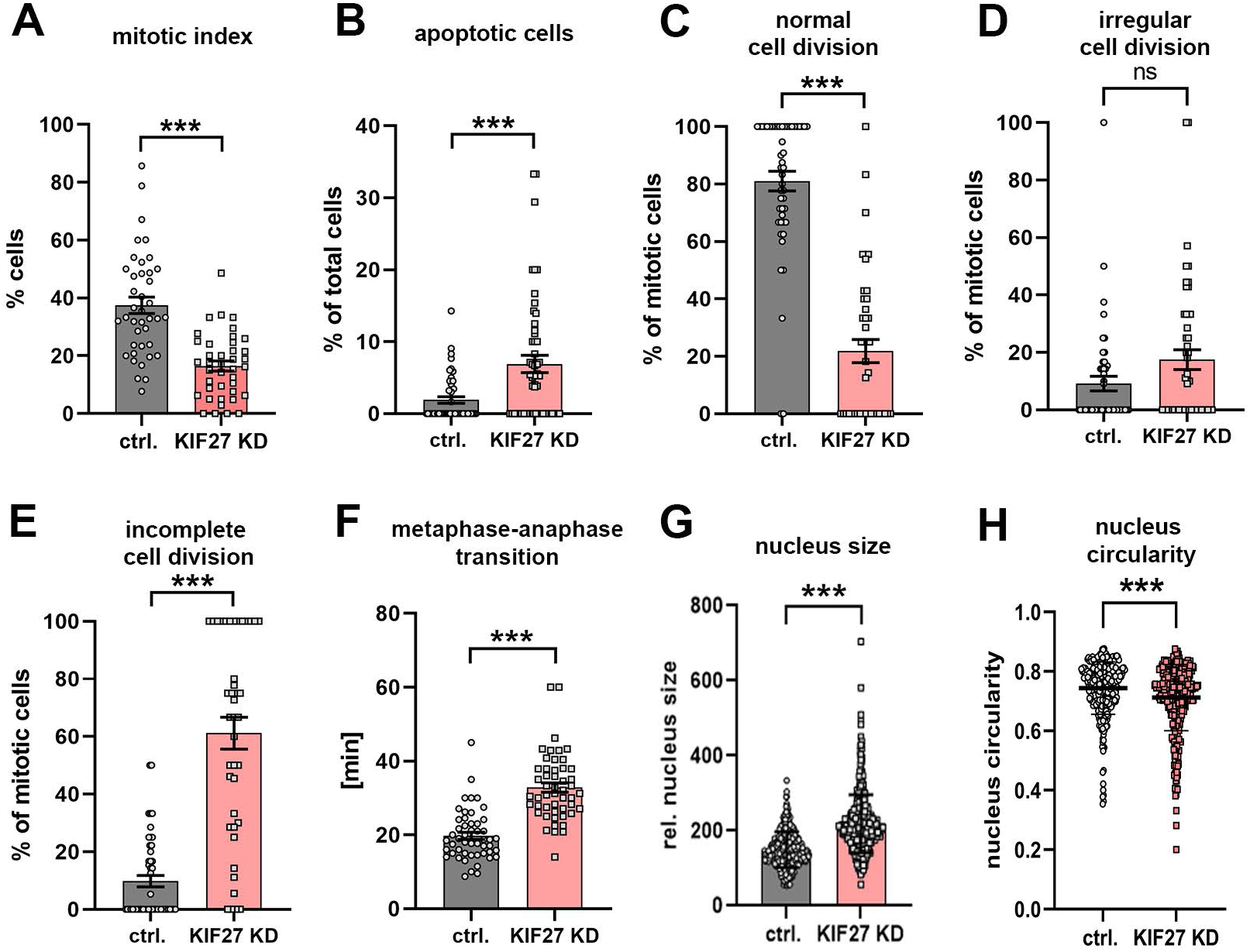
KIF27 depletion causes severe mitotic and cytokinetic defects and interferes with nuclear morphology. Quantitative analysis of cells stained with SiR-tubulin and analyzed by live-cell imaging **(A-F)** or immunofluorescence microscopy of fixed cells stained for tubulin, RacGAP1 and KIF27 **(A)** Scatter plot showing a significantly reduced mitotic index (percentage of total cells) from 37.49 ± 2.56% (ctrl.) to 16.44 ± 2.27% (KIF27 KD) upon KIF27 depletion compared to control cells. **(B)** Significantly increased percentage of apoptotic cells (percentage of total cells) upon KIF27 depletion compared to control cells, from 1.91 ± 0.45% (ctrl.) to 7.05 ± 1.19% (KIF27 KD). **(C)** KIF27 KD significantly decreases the rate of normal cell division with a complete mitotic cycle (percentage of total cells in mitosis) from 81.06 ± 3.44 (ctrl.) to 21.84 ± 4.06% (KIF27 KD). **(D)** KIF27 does not significantly induce irregular cell division compared to control cells (appearance of multipolar spindle structures or multinucleation). Non-significant (ns, P = 0.0675) increase from 9.15 ± 2% (ctrl.) to 16.98 ± 3.45% (KIF27 KD) of the total number of mitotic cells. **(E)** Percentage incomplete cell division (cell cycle arrest, cleavage furrow regression, no abscission of the intercellular bridge) significantly increases after KIF27 depletion compared to control cells, from 9.79 ± 2% (ctrl.) to 61.18 ± 5.52% (KIF27 KD), compared to the sum of all mitotic cells. n >1000 cells from four independent experiments, *P* <0.0001 **(A-E)**. **(F)** Significantly prolonged metaphase-to-anaphase transition time comparing control and KIF27-depleted cells, from 19.76 ± 0.96 min (ctrl.) to 32.81 ± 1.27 min (KIF27 KD). n > 50 cells from three independent experiments. **(G)** Analysis of nucleus size in fixed control cells vs. KIF27 KD cells. KIF27 depletion results in an increase of the relative nucleus size from 150 ± 2.6 a.u. (ctrl.) to 218.9 ± 3.95 a.u. (KIF27 KD). n > 300 cells from three independent experiments. **(H)** Quantification of nuclear circularity following fixation and Hoechst staining in control versus KIF27 KD HeLa K cells. KIF27 depletion significantly reduces nuclear circularity, indicating irregular nuclear shape (mean circularity ± SEM; ctrl.: 0.746 ± 0.005, KD: 0.711 ± 0.006). n > 300 nuclei per condition from three independent experiments. In all graphs error bars indicate ± SEM and *P* values were calculated by unpaired two-sided *t*-test (A-G) or Mann–Whitney U test (H).

### KIF27 regulates the recruitment of midbody-associated proteins and accurate abscission timing

Finally, we addressed whether KIF27 is involved in cytokinetic abscission. Our findings showed that KIF27 depletion can lead to abscission failure (**Fig. 4E**) and irregular midbody morphology (**Fig. 3F-H**). Together with its localization at the midbody and proximity to midbody-associated proteins and abscission factors (**Figures 1**–**3**), these results suggest a role for KIF27 in midbody maturation and abscission. Given that KIF27 was recruited prior to the ESCRT-dependent abscission machinery (ALIX, ESCRT-III/CHMP4B) (**Supplementary Fig. 3A-B**), we analyzed the recruitment of fluorescently labeled CEP55, ALIX and CHMP4B to the midbody in KIF27-depleted versus control cells. KIF27 depletion did not entirely inhibit the midbody recruitment of CEP55, ALIX or CHMP4B, but significantly delayed their appearance at the midbody (**Fig. 5A-C**). We also quantified abscission timing in live-cell imaging experiments of control and KIF27-depleted cells by measuring the interval from formation of a stable cytokinetic bridge to severing of the bridge microtubules (**Fig. 5D**). KIF27-depleted cells exhibited significantly prolonged time to complete abscission compared to control cells (**Fig. 5D**, *P* = 0.0039), in line with the delayed midbody recruitment of CEP55, ALIX and CHMP4B (**Fig. 5A-C**).

**Figure 5.**
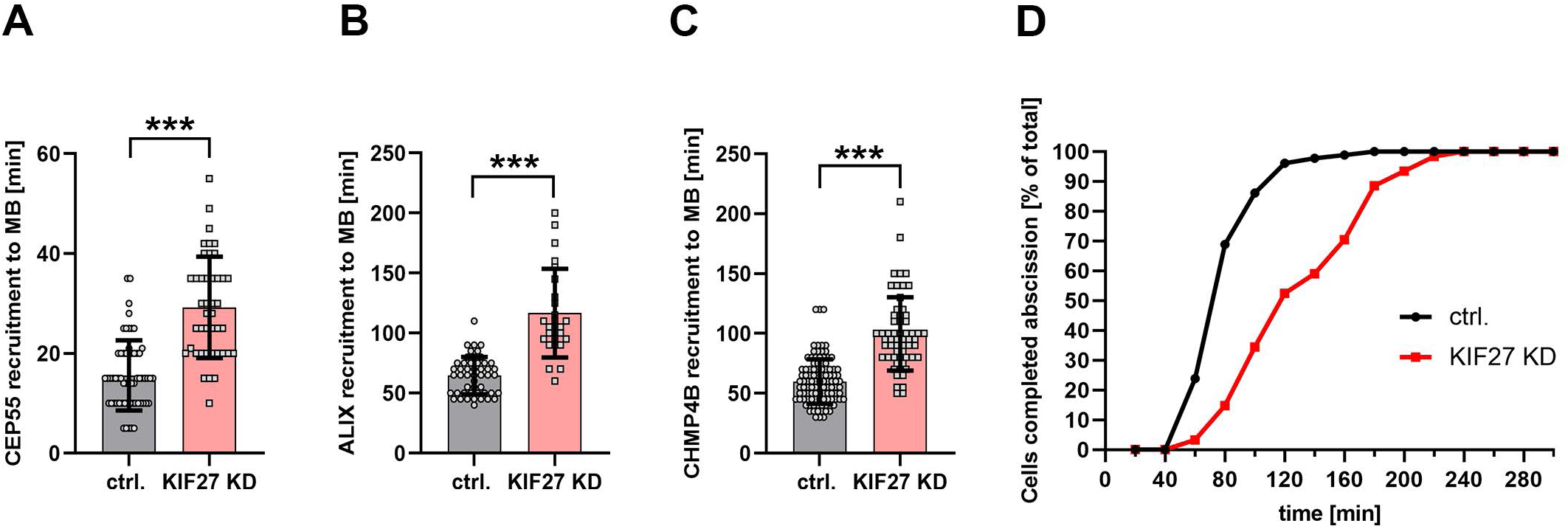
KIF27 regulates the recruitment of midbody-associated proteins and abscission timing. Quantitative analysis of the protein recruitment to the midbody and time of abscission. Data are derived from live-cell imaging experiments comparing control HeLa K cells (ctrl.) and KIF27- depleted HeLa K cells (KIF27 KD). **(A)** Scatter plot showing the time interval between intercellular bridge formation and first appearance of CEP55-mCherry at the midbody (MB) in control cells or upon KIF27 siRNA treatment as indicated (ctrl.: 15.59 ± 0.91 min, KIF27 KD: 29.2 ±1.55 min, n > 40 cells). **(B)** Scatter plot showing the time interval between bridge formation and first appearance of ALIX-mCherry at the midbody (MB) in control cells or upon KIF27 siRNA (KD) treatment as indicated (ctrl.: 64.59 ± 2.23 min, KIF27 KD: 116.7 ± 7.55 min, n > 30 cells). **(C)** Scatter plot showing the time interval between bridge formation and first appearance of CHMP4B-EGFP at the midbody (MB) in control cells or upon KIF27 siRNA (KD) treatment as indicated (ctrl.: 59.92 ± 1.81 min, KIF27 KD: 99.75 ± 3.99 min, n > 60 cells). In all graphs **(A-C)**, error bars indicate ± SEM, *P* < 0.0001, unpaired two-sided *t*-test, three independent experiments. **(D)** Cumulative frequency plot showing the time interval between intercellular bridge formation and abscission in control cells or upon KIF27 depletion. Depletion of KIF27 resulted in a significantly prolonged time to complete abscission (mean time 50% of cells completed abscission ±SEM; ctrl.: 70.35 ± 1.24 min, KD: 124.4 ± 7.4 min). \*\**P* < 0.0039, Wilcoxon signed rank test, n > 80 cells from three independent experiments.

In summary, these results show that KIF27 regulates cytokinesis by coordinating the temporal transition from centralspindlin-driven midzone formation to midbody maturation and ESCRT-III-mediated scission.

### Low KIF27 expression correlates with aneuploidy and decreased patient survival in several cancer types

Given the mitotic defects and nuclear abnormalities caused by KIF27 depletion, we investigated whether KIF27 expression is associated with chromosomal instability in human cancers. We analyzed 9684 tumors across 33 cancer types from The Cancer Genome Atlas (TCGA) and found a significant negative Spearman correlation between KIF27 mRNA expression and aneuploidy score (**Fig. 6A** and **Supplementary Table S1**). Six cancer types reached individual significance after Benjamini-Hochberg false discovery rate (FDR) correction: uterine corpus endometrial carcinoma (UCEC), cervical squamous cell carcinoma (CESC), kidney renal clear cell carcinoma (KIRC), liver hepatocellular carcinoma (LIHC), thyroid carcinoma (THCA), and breast invasive carcinoma (BRCA). In each case, tumors with low KIF27 expression had higher aneuploidy scores (**Fig. 6A** and **Supplementary Fig. 4A-G**). Consistently, KIF27-low tumors also showed broadly elevated chromosome-arm loss frequencies across the genome (**Supplementary Fig. 5A**).

**Figure 6.**
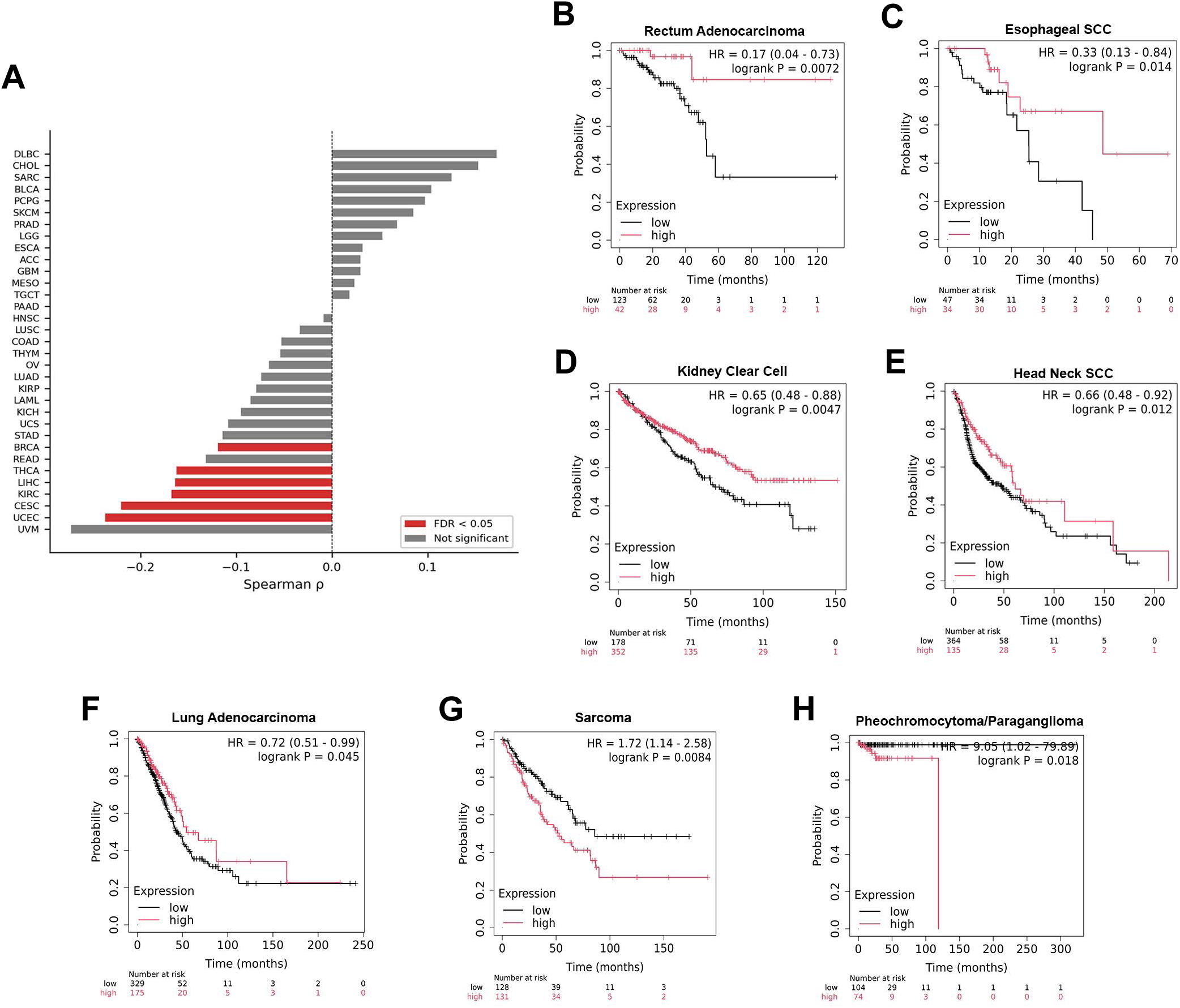
KIF27 expression correlates with aneuploidy and overall survival across TCGA cancer types. **(A)** Forest plot of Spearman correlation coefficients (ρ) between KIF27 mRNA expression and aneuploidy score (Taylor *et al*. 2018) across 33 TCGA cancer types (n = 9,684 tumors), ordered by ρ. Red bars indicate cancer types with statistically significant correlations (FDR < 0.05); grey bars indicate cancer types that are not significant. Negative ρ indicates that higher KIF27 expression is associated with lower aneuploidy. Six cancer types reach individual significance (UCEC, CESC, KIRC, LIHC, THCA and BRCA). See the main text for details. **(B-H)** Individual Kaplan–Meier overall survival curves for selected cancer types with statistically significant correlation between overall survival and KIF27 expression. Survival data from KMplot.com (Győrffy 2024), RNA-seq platform, auto-selected best cutoff. Number at risk is shown below each curve. See the main text and Method section for details. **(B)** Rectum adenocarcinoma: HR = 0.17 (95% CI 0.04-0.73), log-rank p = 0.0072. **(C)** Esophageal squamous cell carcinoma: HR = 0.33 (0.13-0.84), p = 0.014. **(D)** Kidney renal clear cell carcinoma: HR = 0.65 (0.48-0.88), p = 0.0047. **(E)** Head and neck squamous cell carcinoma: HR = 0.66 (0.48-0.92), p = 0.012. **(F)** Lung adenocarcinoma: HR = 0.72 (0.51-0.99), p = 0.045. **(G)** Sarcoma: HR = 1.72 (1.14-2.58), p = 0.0084. **(H)** Pheochromocytoma and paragangliom: HR = 9.05 (1.02-79.89), p = 0.018. Unfavorable; note very wide CI due to n = 7 events.

As KIF27 is located on chromosome 9p21.2, we tested whether the correlation between KIF27 expression and aneuploidy could be explained by cis-dosage effects, in which deletion of the 9p21.2 locus would simultaneously lower KIF27 expression and raise the aneuploidy score. After excluding tumors with 9p deletions, the pan-cancer correlation was retained at 93% of its original magnitude, and five of the six individually significant cancer types remained significant (**Supplementary Fig. 5B-C**). Thus, this correlation is not solely driven by 9p cis-deletion.

We then examined whether KIF27 expression is associated with overall survival. Using Kaplan-Meier plotter, we identified seven cancer types with significant associations between KIF27 expression and overall survival (**Fig. 6B-H**). High KIF27 expression was associated with favorable outcome in rectum adenocarcinoma (HR = 0.17), esophageal squamous cell carcinoma (SCC) (HR = 0.33), KIRC (HR = 0.65), head and neck squamous cell carcinoma (HNSC) (HR = 0.66), and lung adenocarcinoma (LUAD) (HR = 0.72), whereas high KIF27 was associated with unfavorable outcome in sarcoma (HR = 1.72) and pheochromocytoma/paraganglioma (PCPG) (HR = 9.05). These data show that KIF27 expression is associated with overall survival in multiple cancer types, with the direction depending on tumor lineage.

Next, we asked whether the prognostic effect of KIF27 is mechanistically linked to its correlation with aneuploidy. We performed univariate Cox regression of aneuploidy score against overall survival in each of the seven cancer types in which KIF27 expression significantly correlated with survival (**Supplementary Table S2**) and only sarcoma showed a significant aneuploidy to survival association (HR = 1.029 per aneuploidy score unit, P = 0.018) (**Supplementary Table S2**). Moreover, KIRC was the only cancer type where KIF27 expression significantly correlated with both aneuploidy and overall survival, with no correlation for the other six cancer types (**Fig. 6B-H** and **Supplementary Fig. 4A-G**), indicating that the prognostic effect of KIF27 is mechanistically independent of aneuploidy and is therefore likely linked to other pathways.

Taken together, these results indicate that KIF27 expression correlates with chromosomal stability and that KIF27 expression is associated with overall survival in multiple cancer types.

## Discussion

We identify KIF27, a kinesin-4 family member and paralogue of KIF7, as a regulator of mitotic progression and cytokinetic abscission that links the central spindle midzone to timely scission. Although KIF7 and KIF27 are paralogues and both orthologous to *Drosophila* Costal-2 (Park et al., 2025), our data reveal distinct roles during cell division. KIF27 localizes to the central spindle and midbody, whereas KIF7 is not detected at these structures (**Table 1** and **Supplementary Fig. 2H**), demonstrating a mitosis-specific function unique to KIF27.

The dynamic spatiotemporal redistribution that KIF27 exhibits throughout the cell cycle (**Fig 1 A-C**), suggests multiple functions in interphase and during cell division. The mitotic localization partially resembles that of KIF4A (**Fig. 2A**), a known chromokinesin and modulator of chromatin architecture in interphase (Lee et al., 2001), bringing up the possibility that KIF27 may have similar nuclear functions. However, KIF27 shows a less pronounced association with condensed chromosomes compared to KIF4A (**Fig. 2A**), lacks canonical DNA-binding regions, and does not colocalize with KIF4A on chromosomes (**Supplementary Fig. 3E**), suggesting a mechanistically distinct role. In interphase nuclei, nucleoli are sites of ribosomal RNA synthesis and ribosome biogenesis and act as organizing hubs for heterochromatin and nuclear architecture (Boisvert et al., 2007; Iarovaia et al., 2019). The nucleolar localization of KIF27 (**Supplementary Fig. 1A**), suggests a functional role for KIF27 in these structures. In the nucleus, KIF27 also colocalized with RacGAP1 and STAT3 (**Fig. 1B, Supplementary Fig. 1C**). Since RacGAP1 promotes the nuclear translocation of activated STAT transcription factors (Kawashima et al., 2006), KIF27 might contribute to STAT-driven gene expression. These observations suggest a nuclear role of KIF27 similar to RacGAP1; however, its specific interphase functions remain to be determined.

We identified that KIF27 accumulated at the spindle midzone together with KIF4A and the centralspindlin complex components RacGAP1 and KIF23/MKLP1 during anaphase (**Fig. 2A-C**, **Table 1**). The centralspindlin complex is essential for bundling antiparallel microtubules during central spindle formation, which drives contractile ring formation and cleavage furrow ingression (Davies et al., 2015; Fesquet et al., 2015; Maruyama et al., 2021). Disruption of midzone organization, by depletion of KIF4A, RacGAP1 or KIF23, leads to disruption of the central spindle organization, chromosome alignment, cleavage furrow formation and cytokinesis, leading to genomic instability and aneuploidy (Kurasawa et al., 2004; Mazumdar et al., 2006; Mazumdar et al., 2004; Ramkumar et al., 2021). Similarly, KIF27 depletion led to delayed metaphase-anaphase transitions, chromosome missegregation, and defective cytokinesis (**Fig. 4**). These phenotypes align mechanistically with disruption of central spindle microtubule-bundling and protein complex assembly (White and Glotzer, 2012), suggesting that KIF27 has a role in stabilizing the spindle midzone.

Our findings identify KIF27 as a midbody core protein during late stage of cytokinesis. Specifically, KIF27 co-localized with KIF23, PRC1 and CEP55 at the midbody core (**Fig. 2B, C, G** and **Supplementary Fig. 2E**). Moreover, KIF23 and CEP55 promoted KIF27 recruitment (**Fig. 3A-B**), and KIF27 in turn stabilized the midbody to ensure timely recruitment of ALIX and CHMP4B and accurate abscission (**Fig. 5**). The centralspindlin complex thus appears to be upstream of KIF27, which in turn is upstream of the ESCRT machinery during abscission. Importantly, KIF27 depletion also impaired midbody morphology and microtubule organization (**Fig. 3F-H**), placing KIF27 as a midbody core protein required for structural integrity and abscission timing. KIF27 depletion also reduced mitotic activity (**Fig. 4A**) and caused abnormal nuclear sizes (**Fig. 4G**), consistent with the G2 arrest reported for KIF4A, KIF23, and RacGAP1 depletion (Cho et al., 2019; Fischer et al., 2016; Liu et al., 2020). Together, our data thus favor a model in which KIF27 does not function as a conventional motor protein for cargo transport but instead acts as a scaffold protein, stabilizing protein complexes and organizing microtubules, vital for proper central spindle and midbody formation during cell division. This is consistent with recent studies showing that KIF27 has limited motility and scaffolds microtubules in motile cilia (Park et al., 2025; Yue et al., 2018).

Our pan-cancer analyses link low KIF27 expression with aneuploidy in several tumor types (**Fig. 6A, Supplementary Table S1**), independent of a cis-dosage effect (**Supplementary Fig. 5B-C**). This is mechanistically consistent with KIF27 depletion causing compromised ESCRT-III recruitment and cytokinesis failure (**Fig. 4**–**5**), which is a well-established pathway to tetraploidization and aneuploidy (Fujiwara et al., 2005; Ganem et al., 2007). The abscission checkpoint that prevents tetraploidization upon chromosome mis-segregation depends on proper midbody function (Lens and Medema, 2019; Steigemann et al., 2009), and the defects observed upon KIF27 depletion would be expected to compromise this checkpoint. All the six tumors with low KIF27 expression and high aneuploidy are carcinomas (UCEC, CESC, KIRC, LIHC, THCA, and BRCA), originating from epithelial cells, which is in line with our findings in epithelial HeLa K cells *in vitro*. KIF27 was also significantly associated with overall survival in seven cancer types (**Fig. 6B-H**), with high expression being favorable in five types of carcinomas (rectum adenocarcinoma, esophageal SCC, KIRC, HNSC and LUAD), and unfavorable in sarcoma and PCPG. This prognostic behavior differs from that of other cytokinetic kinesins (KIF20A, KIF14 and KIF23), which are typically overexpressed in tumors and correlate with poor outcome and proliferative activity (Cho et al., 2019; Li et al., 2020; Liu et al., 2020), supporting a guardian-type role for KIF27 in maintaining cell division fidelity rather than driving proliferation. However, the survival associations are largely independent of the aneuploidy correlations (**Supplementary Table S2**). This suggests that KIF27’s prognostic value reflects aneuploidy-independent functions, potentially in nuclear organization, midbody integrity, or abscission fidelity. Notably, the unfavorable associations observed in sarcoma and PCPG underscore lineage-specific functions, consistent with the context-dependent nature of aneuploidy in cancer (Ben-David and Amon, 2020; Vasudevan et al., 2021). The observed enlarged nuclei (**Fig. 4G**) and reduced nuclear circularity upon KIF27 depletion (**Fig. 4H**), can functionally be linked to the nuclear and nucleolar localization of KIF27 in interphase cells (**Fig. 1A-B**). Nuclear enlargement fits with a delay in interphase progression, and the loss of circularity suggests that KIF27 helps to maintain nuclear shape. Both phenotypes are common in chromosomally unstable and aneuploid cells (Hosea et al., 2024; Mazzagatti et al., 2024; Smith et al., 2018), which is in line with the elevated aneuploidy of KIF27-low tumors in our pan-cancer analysis (**Fig. 6**).

In conclusion, we identify KIF27 as a novel regulator of mitotic progression, cytokinetic abscission and nuclear morphology. Consistent with this role, low KIF27 expression is associated with aneuploidy in several human cancers, positioning KIF27 as a multifunctional regulator of cell division fidelity and genome stability.

## Materials and Methods

### Cell Culture and Transfections

Human HeLa ‘‘Kyoto’’ (HeLa K), U2OS, PC3, and RPE1 cells were maintained in DMEM (GIBCO) supplemented with 10% FBS, 100U/ml penicillin, and 100mg/ml streptomycin at 37°C under 5% CO2. Stable cell lines expressing fluorescently labelled ALIX or CHMP4B were generated as described elsewhere (Christ et al., 2016; Radulovic et al., 2018). For transient transfection cells were incubated with a mixture of FuGene 6 (Promega) and the plasmid of interest, using a ratio of 3:1 of Fugene 6 to DNA and incubated for 24-48 h. In most cases, cells were transfected in MatTek 3.5 cm dishes using 6 µl of Fugene 6 and 2 µg of DNA. Before cells were used for further analysis the medium was exchanged. The plasmid expressing CEP55-mCherry was kindly provided by Dr. Kay Schink.

### Antibodies and reagents

The following primary antibodies were used for immunoblotting (IB) and immunofluorescence (IF): anti-Anillin (Abcam #ab154337), anti-ALIX and anti-CHMP4B (IB:(Christ et al., 2016)) and anti-ALIX for immunofluorescence (Bio Legend #634502), anti-BAF57 (Abnova #NB-100-2591), anti-Aurora B (Abcam #ab3609), anti-CENP-E (Abcam #ab5093), anti-CEP55 (Abnova #H00055165), anti-Citron Rho-interacting kinase (CITK, BD Transduction Laboratories #611377), anti-KIF11 (Abcam #ab51976), anti-KIF27 (IF: Novus Biologicals, #NBP3-10310; IB: Santa Cruz #sc-390766), anti-KIF23 (IF: Sigma-Aldrich #HPA045208; IB: Abnova # H00009493), anti-KIF4A (IF: Santa Cruz #sc-365145; IB: Abcam #ab38115), anti-KIF7 (Invitrogen #MA5-24382), anti-LaminB1 (Abcam #ab16048), anti-Histone H2B (Abcam #52484), anti-PRC1 (Santa Cruz #sc-9342), anti-RACGAP1 (Abcam #ab2270), and anti-tubulin (Sigma-Aldrich #T5168). Secondary antibodies included anti-mouse, anti-rabbit, and anti-goat Alexa Fluor 488 (Jackson ImmunoResearch), Alexa Fluor 555 (Molecular Probes), Alexa Fluor 568 (Molecular Probes), Alexa Fluor 647 (Jackson ImmunoResearch), and DyLight649 (Jackson ImmunoResearch). Methanol-free 16% paraformaldehyde (PFA) was from Thermo Fisher Scientific, SiR700-tubulin from Spirochrome, Hoechst 33342 from Thermo Fisher Scientific, and nocodazole was purchased from Merck.

### siRNA transfections

Silencer Select siRNAs against ALIX (5′-GCAGUGAGGUUGUAAAUGU-3′), KIF23 (5’-CCUUUGCCGUCAUGCGAAA-3’),CEP55 (5’-GGAGAAGAAUGCUUAUCAAU-3’), KIF27 siRNA sequences: #1: 5’-GCAAGCUGCUUAACGAUAU-3’, #2: 5’-GGUUCAAGCUGCUUAACGA-3’, #3: 5’-GCAACACAAACUGCAGGAGUGGUUU-3’, #4: 5’-AAACCACUCCUGCAGUUUGUGUUGC -3’) and non-targeting control siRNA (Silencer Select Negative Control No.1 siRNA Cat #4390843) were purchased from Thermo Fisher Scientific. Cells were seeded in six-well plates at 30% confluence and transfected with 12.5-25 nM final siRNA concentration using Lipofectamine RNAiMax (Life Technologies) according to the manufacturers’ instructions and cells were then used for experiments 48 h after knockdown. Knockdown cells used in the experiments showed ≥80% reduced protein levels compared to control cells, as determined by Western blot analysis. For the knockdown of KIF27 we tested transiently transfected four different RNAi oligos. All oligos reduced the KIF27 protein levels to 50-80% (**Supplementary Fig. 3C**) and showed, depending on the efficacy of protein depletion, similar effects in experimental settings. KIF27 knockdown experiments shown in the manuscript were performed with oligo #3.

### Protein extraction and cell fractionation

Proteins were isolated in 3 cellular fractions (cytosolic, nuclear, chromatin bound) as described elsewhere (Mallet et al., 2016). Briefly, cells were resuspended for 30 min at 4°C in cytosolic lysis buffer (10 mM Tris pH 8, 0.34 M sucrose, 3 mM CaCl_2_, 2 mM MgCl_2_, 0.1 mM EDTA, 1 mM DTT, 0.5% NP40) with 40 μl/mL of protease inhibitor (Complete EDTA free, Roche) and centrifuged (5 000 rpm, 2 min). The supernatant was kept as the cytosolic fraction. Pellets containing nuclear and chromatin bound fractions were resuspended in 2.5 vol of nuclear lysis buffer (20 mM HEPES pH 7.9, 1.5 mM MgCl_2_, 1 mM EDTA, 150 mM KCl, 0.1% NP40, 1 mM DTT, 10% glycerol with 40 μl/mL of protease inhibitor). Nuclei were broken by multiple pipetting through a needle (21G) and centrifugation (14,000 rpm, 30 min). The supernatant was isolated as the nuclear fraction. The residual pellet containing the chromatin fraction was resuspended for 1 h, at 4 °C in 2 vol of nuclease incubation buffer (20 mM HEPES pH 7.9, 1.5 mM MgCl_2_, 150 mM KCl, 10% glycerol and 0.15 units/μl benzonase) and centrifuged (14,000 rpm, 3 min) to remove debris. The supernatant was isolated as the chromatin fraction.

### Immunoblotting

Cells were washed with ice-cold PBS and lysed in 2 × sample buffer (125 nM Tris–HCl, pH 6.8, 4% SDS, 20% glycerol, 200 nM DTT and 0.004% bromophenol blue). Wholecell lysates were subjected to SDS–PAGE on 4–20% gradient gels (Mini-PROTEAN TGX; Bio-Rad). Proteins were transferred to polyvinylidene difluoride (PVDF) membranes (Trans-Blot®Turbo™LF PVDF, Bio-Rad) followed by blocking in 5% BSA/PBS and primary antibody incubation in 5% BSA in Tris-buffered saline with 0.1% Tween-20 overnight at 4 °C. Following three washes with PBS/0.01% Tween-20, the membranes were incubated for 45 min with the fluorescent secondary antibodies IRDye680 or IRDye800 (LI-COR, 926–32212, 926–68073, 1:10,000 in 5% BSA/PBS), washed twice in PBS/0.01% Tween-20 and once in PBS, followed by scanning using an Odyssey infrared scanner (LI-COR). Quantification of immunoblots was performed using ImageJ/FIJI.

### Co-immunoprecipitation

For immunoprecipitation experiments KIF27, KIF23, RacGAP1, or CEP55 antibodies (described above) were conjugated with Dynabeads® Protein A or G, as described in the manufacturer’s instructions (Thermo Fisher Scientific), unconjungated Dynabeads were used as a negative control for IP. Cells were seeded the day before, resuspended for 15 min at 4°C in lysis buffer (20 mM HEPES pH 7.9, 1.5 mM MgCl_2_, 1 mM EDTA, 150 mM KCl, 0.1% NP40, 1 mM DTT, 10% glycerol with 40 μl/mL of protease inhibitor). Nuclei were broken by multiple pipetting through a needle (21G) and centrifugation (14,000 rpm, 30 min). The supernatant was incubated for 15 min at room temperature (RT) with unconjugated or antibody-conjungated Dynabeads. After washing and processing of the probes as described in the manufacturer’s protocol, the samples were analyzed by SDS-Page and conjugates were probed with specific antibodies against KIF27, KIF23, RacGAP1, or CEP55.

### Fluorescence microscopy

For immunofluorescence, cells grown on coverslips were fixed in 4% paraformaldehyde for 15 min, permeabilized for 5 min with 0.1% Triton X-100/PBS, and blocked in 10% FBS/PBS. Primary antibodies were applied at RT for 60-75 min, followed by 40 min incubation with appropriate fluorophore-conjugated secondary antibodies (Jackson ImmunoResearch) diluted in PBS/Tween at RT. DNA was stained with Hoechst 33342 (1:2000). Coverslips were mounted in ProLong Glas antifade medium. Images were deconvolved using NIS Elements AR software and processed with FIJI/ImageJ (Schindelin et al., 2012), Icy (de Chaumont et al., 2012), and CellProfiler (Stirling et al., 2021) software.

### Live-cell microscopy

For long-term live-cell microscopy (12-16 h; analysis of cell abscission, protein recruitment during cytokinesis) we used either a DeltaVision microscope (Applied Precision) equipped with an Elite TruLight Illumination System, a CoolSNAP HQ2 camera and a 60× Plan Apochromat (1.42 numerical aperture, NA) lens or a Nikon ECLIPSE Ti2-E inverted microscope equipped with a CrestOptics X-Light V3 spinning disk confocal module, a dual camera imaging setup (back-illuminated sCMOS cameras, Photometrics Kinetix) and a CFI Plan Apo 100X Oil *NA 1.45* lens. For temperature control during live observation, the microscope stage was kept at 37°C by a temperature-controlled incubation chamber. Cells were imaged in live-cell imaging buffer (Invitrogen, supplemented with FBS, penicillin/streptomycin, HEPES and Aln-Gln) at 37°C. Time-lapse images were deconvolved using softWoRx software (Applied Precision, GE Healthcare) or NIS Elements AR software and processed with FIJI/ImageJ (Schindelin et al., 2012).

### Structured illumination microscopy (SIM)

For 3D-SIM (structured illumination microscopy) cells were seeded on coverslips and fixed in 4% EM-grade paraformaldehyde for 15 min (Scheffler et al., 2014) and permeabilized with 0.1% Triton X-100 in PBS for 5min. For protein staining, specific primary antibodies were used as described above. Coverslips were mounted in ProLong™ Gold or ProLong™ Glass (Thermo Fisher Scientific). 3D-SIM imaging was performed on a Deltavision OMX V4 system (Applied Precision) equipped with an Olympus 60x numerical aperture (NA) 1.42 objective, and three PCO.edge sCMOS cameras and 405, 488, 568 and 642 nm diode lasers. Z-stacks covering the whole cell were recorded with a Z-spacing of 125 nm. A total of 15 raw images (five phases, three rotations) per plane were collected and reconstructed by using SOFTWORX software (Applied Precision) and processed in FIJI/ImageJ (Schindelin et al., 2012) and three-dimensional reconstruction was calculated and visualized by icy imaging software (de Chaumont et al., 2012).

### STED nanoscopy

For STED imaging cells were fixed, permeabilized and stained with primary antibodies as described above (Fluorescent microscopy). Afterwards, cells were incubated for 40 min with STED compatible fluorophore-conjugated secondary antibodies (abberior STAR RED and STAR ORANGE, Abberior) diluted in PBS/Tween at RT. STED imaging was performed using an Abberior Instruments Expert Line system operated in a time-gated configuration and mounted on an inverted Olympus IX83 microscope. Images were acquired with an Olympus UPlanSApo 100×/1.4 NA objective using immersion oil with a refractive index of 1.518. Abberior STAR ORANGE and abberior STAR RED were excited with 561- and 640-nm laser lines, respectively, and a 775-nm STED beam was used for depletion in both channels. Fluorescence detection was performed in the 595–665 nm range for the abberior STAR ORANGE and in the 650–750 nm range for abberior STAR RED. For each region of interest, Z-stacks spanning several micrometers (depending on the size of the imaged structures) were acquired with a z-step size of 50 nm. For deconvolution, xy scanning was oversampled using a pixel size of 10 nm. STED images were deconvolved using Huygens Professional software (SVI, The Netherlands) with the express wizard in standard profile, applying a STED saturation factor of 30 and a STED immunity fraction of 10%.

### Proximity Ligation Assay (PLA)

The Duolink® PLA (Merck) was used to detect close proximity of endogenous KIF27 with ALIX, CENP-E, and Aurora B. The assay is based on oligonucleotide-conjugated PLA probes, containing secondary antibodies directed against primary antibodies against the proteins of interest. Annealing of the probes occurs when the target proteins are in close proximity (<40 nm), which then initiates the amplification. The amplicons can be detected by fluorescence microscopy in a quantifiable manner. For this assay, Hela K cells were seeded on coverslips in six-well plates and the PLA experiments were performed the next day with subconfluent cells. Cells were washed, fixed and permeabilized as described in the section above. Antibodies against KIF27 (1:100, Novus Biologicals, #NBP3-10310), ALIX (1:100, BioLegend #634502), CENP-E (1:100, Abcam #ab5093), and Aurora B (1:100, Abcam #ab3609) were used as primary antibodies, and the assays were performed as described in the manufacturer’s manual. Slides were mounted with ProLong Glass (Invitrogen) and samples were observed by fluorescence microscopy with a Nikon ECLIPSE Ti2 spinning-disk microscope. PLA signals were visualized by confocal microscopy and quantified using FIJI/ImageJ (Schindelin et al., 2012).

### STEM tomography

Cells were grown on coverslips and then fixed with 2% glutaraldehyde in 0.1 M PHEM buffer (60 mM PIPES, 25 mM HEPES, 2 mM MgCl2, 10 mM EGTA, pH 6.9) for 1 h. Postfixation was done in 1% OsO4 and 1.5% KFeCN in the same buffer. Samples were further stained en bloc with 4% aqueous uranyl acetate for 1 h, dehydrated in graded ethanol series and embedded with Epon-filled gelatine capsules (EMS Polysciences Inc.) placed on top of the coverslip. After polymerization serial sections (750 nm) were cut on an Ultracut UCT ultramicrotome (Leica, Germany) and collected on formvar-coated slot grids. Samples were observed in a Thermo Fisher Scientific™ Talos™ F200C microscope at 200 kV using the bright field detector for STEM (scanning transmission electron microscopy) imaging with microprobe mode. The convergence angle of the scanning beam was 1.7 mrad, condenser aperture of 70 nm and a camera length of 530 mm. For STEM tomography image series were taken at - 56° to 56° tilt angles with 2° increment and a pixel size of 2.25 nm. Tomograms were computed using weighted back projection using the IMOD package. Display of tomogram slices was also performed using IMOD software version 4.9.3

### TCGA expression, aneuploidy and survival analysis

Gene-level RNA-seq expression data (EB++AdjustPANCAN, IlluminaHiSeq) were obtained from the TCGA Pan-Cancer Atlas. This matrix is the Pan-Cancer Atlas expression dataset in which cross-tumor batch and platform effects had been corrected by the consortium using an Empirical-Bayes (ComBat-type) adjustment, denoted “EB++”. Aneuploidy scores were taken from Taylor *et al*. (2018). Spearman rank correlations between KIF27 expression and aneuploidy score were computed per cancer type and pan-cancer, with Benjamini–Hochberg FDR correction for 33 comparisons. Sensitivity analyses excluded tumors with chromosome 9p deletions to control for potential cis-dosage effects on KIF27 expression at 9p21.2. KIF27 expression-survival associations were analyzed using KMplot (kmplot.com; Győrffy 2024) across 21 RNA-seq cancer types with auto-selected best cutoff. Hazard ratios (HR) express the relative risk of death for patients with high versus low KIF27 expression (or per one-unit increase of a continuous covariate); an HR below 1 indicates a lower risk (favorable prognosis), whereas an HR above 1 indicates a higher risk (unfavorable prognosis). The 95% confidence interval (CI) indicates the range in which the true hazard ratio is expected to lie with 95% confidence, and a CI that excludes 1 denotes a statistically significant association. The log-rank p-value tests the null hypothesis of no difference in overall survival between the high- and low-KIF27 expression groups, with a value below 0.05 considered statistically significant. The Kaplan-Meier plot survival analyses were based on the internally normalised RNA-seq database provided by KMplot and are therefore independent of the EB++/z-scored expression values used for the correlation and multivariate Cox analyses. Aneuploidy–survival associations were computed using overall survival data from the TCGA Pan-Cancer Clinical Data Resource (Liu *et al*. 2018) merged with Taylor *et al*. (2018) aneuploidy scores by patient barcode (n = 9684 matched patients). Univariate Cox proportional-hazards regression of aneuploidy score against overall survival was performed for each of the seven KIF27–survival-significant cancer types. All survival analyses were performed using the lifelines Python package (v0.28); survival time was converted from days to months.

### Statistical Analysis

Statistical analysis was carried out in GraphPad Prism (GraphPad Software). Student’s *t*-test was used to compare two groups. ANOVA was used to compare multiple groups and Holm–Sidak was used to correct for multiple comparisons. The threshold for significance was set at *P* = 0.05. All comparisons made are reported regardless of significance. Comparisons in the Figures are indicated as ns (not significant): *P* ≥ 0.05, *: *P* < 0.05, **: *P* < 0.01, ***: *P* < 0.001.

## Data Availability

All data supporting the conclusions of this article are included within the article and its supplementary files. Additional data and information are available from the corresponding author upon reasonable request.

## Acknowledgments

We thank members of the Stenmark lab for helpful discussions and technical assistance. We acknowledge the Core Facilities for Advanced Light Microscopy and the Advanced Electron Microscopy at the Oslo University Hospital.

## Author Contributions

S.P. and K.H. initiated the project, and all authors contributed to the study conception and design. Material preparation, data collection and analysis were performed by S.P., S.M.M. and A.B. Validation, data curation and visualization were performed by S.P. The original draft of the manuscript was written by S.P. and K.H. All authors reviewed and edited the manuscript. K.H. and H.S. supervised the study. K.H., H.S. and S.G.S. acquired funding.

---

**Supplementary Figure 1. KIF27 association with nucleolin and STAT3, detection of KIF4A in subcellular fractions, and prediction of KIF27 nuclear localization signals.**

**(A)** Immunofluorescence (IF) images showing KIF27 (magenta) localization within the nucleoli in interphase cells, as demonstrated by colocalization with nucleolin (green). DNA is stained with Hoechst (yellow). Scale bar: 10 µm.

**(B)** Western blot analysis of subcellular fractions (cytosol, nucleus, chromatin, and whole cell lysate) confirming the presence of KIF4A in cytoplasmic, nuclear, and chromatin-enriched fractions. Co-immunoprecipitation of can be observed in the cytosolic and nuclear fractions. RacGAP1 and GAPDH was used to monitor fraction purity.

**(C)** Immunofluorescence analysis of interphase HeLa K cells showing partial colocalization of KIF27 (magenta) with STAT3 (green) in the nucleus. Hoechst is stained in blue. Scale bars: 10 µm.

**(D)** Human KIF27 amino acid sequence, with putative nuclear localization sequences (NLS) predicted by cNLS Mapper (https://nls-mapper.iab.keio.ac.jp/cgi-bin/NLS_Mapper_form.cgi) shown in red.

**Supplementary Figure 2. KIF27 localizes to the midbody across cell types and associates with abscission and central-spindle proteins.**

**(A)** IF images show the recruitment of KIF27 (magenta) to the midbody across multiple human cell types (PC3, U2OS and RPE-1). Co-staining with KIF23 (PC3, green) or RacGAP1 (U2OS and RPE-1, green) and tubulin (blue). Scale bars: 2 µm.

**(B)** IF images of HeLa K cell with co-localization of KIF27 (magenta) and KIF23 (green) at the midbody in an intercellular bridge. Tubulin is shown in blue. Scale bar: 1 µm.

**(C)** Detection of endogenous proximity showing the spatial association (≤40 nm) of KIF27 with ALIX, CENP-E, and Aurora B by Duolink® PLA in HeLa K cells. Arrowheads indicate examples of protein proximity at the bridge, represented by fluorescent dots (magenta). Tubulin is shown in blue. Scale bars: 2 µm.

**(D - G)** IF images of HeLa K cells showing KIF27 (magenta) relative localization with different midbody-associated proteins (green): CHMP4B **(D)**, PRC1 **(E)**, Anillin **(F**, SIM), and CITK **(G)**. The microtubuli in the bridge are stained in blue. Scale bars: 1 µm.

**(H + I)** IF images showing that midbody-localized KIF27 (magenta) does not exhibit co-localization with its paralogue KIF7 (green) **(H**, SIM), nor does it show significant colocalization with the cytokinetic motor protein KIF11 (green) **(I,** STED) in HeLa K cells. Tubulin is shown in blue in **(H)**. Scale bars: 1µm.

**Supplementary Figure 3. Characterization of KIF27 localization, knockdown validation, and candidate interaction partners.**

Structured illumination microscopy (SIM) projection images showing the localization of ALIX (white), KIF27 (magenta) and RacGAP1 (green) in fixed **(A)** control HeLa K cells (ctrl.) and **(B)** HeLa K cells depleted of ALIX (ALIX KD). Depletion of ALIX did not prevent recruitment of KIF27 or RacGAP1 to the midbody. Scale bars: 1 µm. See also Movie 3 and 4 for the corresponding 3D SIM animations.

**(C**) Western blot confirming the efficiency of KIF27 depletion using four different KIF27 siRNA oligonucleotides (siKIF27 #1, #2, #3, #4) compared to the control (ctrl.) in HeLa K cells. All oligos reduced KIF27 protein levels by 50–80%. GAPDH serves as a loading control.

**(D)** IF micrographs of nuclei (Hoechst, yellow) in control (ctrl.) and KIF27 siRNA-treated (KD) HeLa K cells. Depletion of KIF27 led to significant increase of the nuclear size (quantification shown in Fig. 4G). Scale bars: 10 µm.

**(E)** Immunofluorescence image showing the comparison of KIF27 (magenta) and KIF4A (green) localization on condensed chromosomes (Hoechst, yellow) in anaphase HeLa K cell. The study revealed no significant colocalization of KIF27 and KIF4A at condensed chromosomes, suggesting distinct roles. Scale bars: 5 µm (left) and 2 µm (insets).

**Supplementary Figure 4. Low KIF27 expression correlates with aneuploidy across cancer types.**

**(A–F)** Box-and-whisker plots comparing aneuploidy score between KIF27-Low (pink) and KIF27-High (grey) expression tertiles in six cancer types with significant KIF27–aneuploidy correlations (FDR < 0.05). Individual data points are shown as open circles. **(A)** Uterine corpus endometrial carcinoma (UCEC; n = 171 vs. 175). **(B)** Cervical squamous cell carcinoma (CESC; n = 98 vs. 102). **(C)** Kidney renal clear cell carcinoma (KIRC; n = 162 vs. 167). **(D)** Liver hepatocellular carcinoma (LIHC; n = 120 vs. 119). **(E)** Thyroid carcinoma (THCA; n = 154 vs. 156). **(F)** Breast invasive carcinoma (BRCA; n = 348 vs. 351). Boxes show 25th–75th percentiles with median line; whiskers extend to 1.5 times the interquartile range (IQR). *, P < 0.05; **, P < 0.01; ***, P < 0.001 (Mann–Whitney U test). Expression and aneuploidy data from TCGA/UCSC Xena and Taylor et al. 2018.

**(G)** Distribution of aneuploidy scores in KIF27-Low (lowest expression tertile, pink) versus KIF27-High (highest expression tertile, blue) tumors across TCGA samples. KIF27-Low tumors show a shift toward higher aneuploidy scores (Mann–Whitney P = 2.45 × 10⁻⁴⁴). Expression data: TCGA Pan-Cancer EB++AdjustPANCAN (UCSC Xena); aneuploidy scores from Taylor et al. *Cancer Cell* 2018. The y-axis (density) is the probability density of the aneuploidy score, that is, the relative frequency of tumors at each score normalized so that the total area under each histogram equals 1. This normalization makes the shapes of the two distributions directly comparable even though the groups differ in size.

**Supplementary Figure 5. Chromosome-arm loss patterns and the contribution of 9p cis-deletion to the KIF27–aneuploidy association.**

**(A)** Horizontal bar plot showing the difference in arm loss frequency (Δ = frequency in KIF27-Low tertile − frequency in KIF27-High tertile) for each of 39 autosomal chromosome arms, ordered by Δ. Positive values indicate preferential arm loss in KIF27-Low tumors. Chromosome 9q and 9p show the largest differential loss, consistent with cis-deletion of the KIF27 locus at 9p21.1. Red bars: FDR < 0.05; grey: not significant.

**(B)** Effect of 9p cis-deletion on the pan-cancer KIF27-aneuploidy correlation. Bars show |Spearman ρ| between KIF27 expression and aneuploidy score for all tumors (n = 9684), tumors excluding those with 9p deletion (n = 7249; 93% of original effect retained), and tumors with 9p deletion only (n = 2435; 34% of original effect). The persistence of the association after excluding 9p-deleted tumors indicates that the KIF27-aneuploidy link is not solely driven by cis-deletion.

**(C)** Per-cancer |Spearman ρ| with (blue) and without (orange) 9p-deleted tumors for the six significant cancer types in Supplementary Figure 4A-F. Asterisks indicate cancer types retaining significance after 9p exclusion. Not significant (n.s.) in LIHC indicates loss of significance after 9p exclusion, suggesting partial cis contribution in this type.

**Supplementary Table S1.**

Per-cancer type Spearman correlations between KIF27 mRNA expression and aneuploidy score in TCGA. Sorted by Spearman correlation coefficient, ρ (most negative first). Significant (Sig.) correlations (FDR < 0.05) in bold. Related to Figure 6A. Each cancer type is shown with its TCGA study abbreviation in parentheses, and significant types are additionally flagged with a check mark (✓) in the Sig. column. Spearman *ρ* ranges from −1 to +1 and measures the strength and direction of the monotonic association between KIF27 mRNA level and aneuploidy score. A negative *ρ* denotes an inverse relationship, meaning that tumors with higher KIF27 expression tend to have a lower aneuploidy score, while a positive *ρ* means that higher KIF27 expression tends to accompany a higher aneuploidy score, and values near 0 indicate no consistent association. *n* is the number of tumor samples analyzed. The *P*-value is the probability of observing a correlation at least this strong if KIF27 expression and aneuploidy score were unrelated, and FDR *q* is that *P*-value after Benjamini–Hochberg adjustment for testing many cancer types. A check mark (✓) marks the types passing the FDR *q* < 0.05 threshold.

**Supplementary Table S2.**

KIF27 mRNA expression and baseline aneuploidy score as predictors of overall survival across seven RNA-seq-significant TCGA cancer types. Shown are seven TCGA cancer types in which KIF27 mRNA reaches significance for overall survival on the Kaplan-Meier plot RNA-seq platform. For each type: sample size (n), events, univariate hazard ratios (HR) and log-rank P-values for KIF27 expression and baseline aneuploidy score (AS; univariate Cox regression, TCGA). Bold values and red shading denote significance (*P < 0.05) in Sarcoma. Events are the number of patients who died during follow-up, that is, the deaths used as the survival endpoint in the analysis. The 95% confidence interval (CI) is the range expected to contain the true hazard ratio with 95% confidence, and an interval that excludes 1 indicates a statistically significant effect. Esoph. SCC, esophageal squamous cell carcinoma; HNSC, head and neck squamous cell carcinoma; KIRC, kidney renal clear cell carcinoma; LUAD, lung adenocarcinoma; PCPG, pheochromocytoma and paraganglioma; Rectum, rectum adenocarcinoma; Sarcoma, soft-tissue sarcoma; OS, overall survival; CI, confidence interval.

**Movie 1**

Localization of KIF27 (magenta) and RacGAP1 (green) at the midbody in fixed cells. Animated projections of 3D reconstructed spinning disk microscopy data from Fig. 2F (middle panel) at different visual angles. Hela K cells were stained for endogenous KIF27, RacGAP1 and tubulin (blue).

**Movie 2**

Morphological changes of KIF27 during the maturation of the intercellular bridge and midbody during cytokinesis. Animated projections of 3D reconstructed SIM images at different visual angles showing the localization of KIF27 (magenta) and ALIX (green) in the intercellular bridge (tubulin, blue) during different progressive stages of cytokinesis (from top to bottom). KIF27 accumulates at the midbody prior to ALIX. Both proteins show a dynamic morphological change during the maturation of the bridge and midbody. See also corresponding Fig. 2G.

**Movie 3**

Localization of ALIX (white), KIF27 (magenta) and RacGAP1 (green) at the midbody of control HeLa K cells. Animated projections of 3D reconstructed SIM data from Suppl. Fig. 3A at different visual angles. Scale bar: 1 µm.

**Movie 4**

Animated projections of 3D reconstructed SIM images showing the localization of ALIX (white), KIF27 (magenta) and RacGAP1 (green) in the intercellular bridge of a ALIX-depleted HeLa K cell. Animated projections of 3D reconstructed SIM data from Suppl. Fig. 3B at different visual angles. Depletion of ALIX does not inhibit the recruitment and localization of KIF27 and RacGAP1 at the midbody. Scale bar: 1 µm.

**Movie 5**

Scanning transmission electron microscopy (STEM) tomogram of an intercellular bridge and midbody of a cytokinetic KIF27-depleted HeLa K cell. The tomogram reveals a loose microtubule system with enlarged vesicular structures on both sides of the midbody and in the intercellular bridge. See also corresponding Fig. 3H.

**Movie 6**

Induction of apoptosis in KIF27-depleted cells. Time-lapse imaging of an interphase Hela K cell upon treatment with KIF27 siRNA and addition of SiR-tubulin during the indicated period in hours and minutes (see also corresponding Fig. 4B). Scale bar: 10 µm.

**Movie 7**

Normal cell division of a Hela K cell. Time-lapse imaging of an interphase cell after addition of SiR-tubulin performing normal mitosis and cytokinesis during the indicated period in hours and minutes (see also corresponding Fig. 4C). Scale bar: 10 µm.

**Movie 8**

Irregular cell division of a Hela K cell. Time-lapse imaging of an interphase cell treatment with KIF27 siRNA and addition of SiR-tubulin during the indicated period in hours and minutes (see also corresponding Fig. 4D). The cell shows formation of three spindle pols, accompanied by an elongated intercellular bridge and delayed abscission. Scale bar: 10 µm.

**Movie 9**

Mitotic arrest of a Hela K cell. Time-lapse imaging of an interphase cell upon treatment with KIF27 siRNA and addition of SiR-tubulin during the indicated period in hours and minutes (see also corresponding Fig. 4E). The cell enters mitosis and arrests in metaphase after the formation of a mitotic spindle with two spindle pols. Scale bar: 10 µm.

**Movie 10**

Delayed meta-to-anaphase transition upon KIF27 depletion. Time-lapse microscopy of DNA-stained (spy650-DNA) control HeLa K cells (left) and HeLa K cells depleted for KIF27 (right) during the indicated period in hours and minutes (see also corresponding Fig. 4F). Depletion of KIF27 leads to a significantly prolonged meta-to-anaphase transition, accompanied by problems with chromosome segregation and the appearance of lagging chromosomes.

## References

Addi, C., A. Presle, S. Fremont, F. Cuvelier, M. Rocancourt, F. Milin, S. Schmutz, J. Chamot-Rooke, T. Douche, M. Duchateau, Q.G. Gianetto, A. Salles, H. Menager, M. Matondo, P. Zimmermann, N. Gupta-Rossi, and A. Echard. 2020. The Flemmingsome reveals an ESCRT-to-membrane coupling via ALIX/syntenin/syndecan-4 required for completion of cytokinesis. Nature Communications. 11.

Agromayor, M., and J. Martin-Serrano. 2013. Knowing when to cut and run: mechanisms that control cytokinetic abscission. Trends Cell Biol. 23:433–441.

Almeida, A.C., and H. Maiato. 2018. Chromokinesins. Curr Biol. 28:R1131–R1135.

Andrade, V., and A. Echard. 2022. Mechanics and regulation of cytokinetic abscission. Front Cell Dev Biol. 10:1046617.

Ben-David, U., and A. Amon. 2020. Context is everything: aneuploidy in cancer. Nat Rev Genet. 21:44–62.

Boisvert, F.M., S. van Koningsbruggen, J. Navascues, and A.I. Lamond. 2007. The multifunctional nucleolus. Nat Rev Mol Cell Biol. 8:574–585.

Carlton, J.G., M. Agromayor, and J. Martin-Serrano. 2008. Differential requirements for Alix and ESCRT-III in cytokinesis and HIV-1 release. Proc Natl Acad Sci U S A. 105:10541–10546.

Carlton, J.G., and J. Martin-Serrano. 2007. Parallels between cytokinesis and retroviral budding: a role for the ESCRT machinery. Science. 316:1908–1912.

Cheung, H.O., X. Zhang, A. Ribeiro, R. Mo, S. Makino, V. Puviindran, K.K. Law, J. Briscoe, and C.C. Hui. 2009. The kinesin protein Kif7 is a critical regulator of Gli transcription factors in mammalian hedgehog signaling. Sci Signal. 2:ra29.

Cho, S.Y., S. Kim, G. Kim, P. Singh, and D.W. Kim. 2019. Integrative analysis of KIF4A, 9, 18A, and 23 and their clinical significance in low-grade glioma and glioblastoma. Sci Rep. 9:4599.

Christ, L., E.M. Wenzel, K. Liestol, C. Raiborg, C. Campsteijn, and H. Stenmark. 2016. ALIX and ESCRT-I/II function as parallel ESCRT-III recruiters in cytokinetic abscission. J Cell Biol. 212:499–513.

D’Avino, P.P., and L. Capalbo. 2016. Regulation of midbody formation and function by mitotic kinases. Semin Cell Dev Biol. 53:57–63.

Davies, T., N. Kodera, G.S. Kaminski Schierle, E. Rees, M. Erdelyi, C.F. Kaminski, T. Ando, and M. Mishima. 2015. CYK4 promotes antiparallel microtubule bundling by optimizing MKLP1 neck conformation. PLoS Biol. 13:e1002121.

de Chaumont, F., S. Dallongeville, N. Chenouard, N. Herve, S. Pop, T. Provoost, V. Meas-Yedid, P. Pankajakshan, T. Lecomte, Y. Le Montagner, T. Lagache, A. Dufour, and J.C. Olivo-Marin. 2012. Icy: an open bioimage informatics platform for extended reproducible research. Nature methods. 9:690–696.

Dong, Z., C. Zhu, Q. Zhan, and W. Jiang. 2018. Cdk phosphorylation licenses Kif4A chromosome localization required for early mitotic progression. J Mol Cell Biol. 10:358–370.

Endoh-Yamagami, S., M. Evangelista, D. Wilson, X. Wen, J.W. Theunissen, K. Phamluong, M. Davis, S.J. Scales, M.J. Solloway, F.J. de Sauvage, and A.S. Peterson. 2009. The mammalian Cos2 homolog Kif7 plays an essential role in modulating Hh signal transduction during development. Curr Biol. 19:1320–1326.

Fabbro, M., B.B. Zhou, M. Takahashi, B. Sarcevic, P. Lal, M.E. Graham, B.G. Gabrielli, P.J. Robinson, E.A. Nigg, Y. Ono, and K.K. Khanna. 2005. Cdk1/Erk2- and Plk1-dependent phosphorylation of a centrosome protein, Cep55, is required for its recruitment to midbody and cytokinesis. Dev Cell. 9:477–488.

Fesquet, D., G. De Bettignies, M. Bellis, J. Espeut, and A. Devault. 2015. Binding of Kif23-iso1/CHO1 to 14-3-3 is regulated by sequential phosphorylations at two LATS kinase consensus sites. PLoS One. 10:e0117857.

Fischer, M., M. Quaas, L. Steiner, and K. Engeland. 2016. The p53-p21-DREAM-CDE/CHR pathway regulates G2/M cell cycle genes. Nucleic Acids Res. 44:164–174.

Fujiwara, T., M. Bandi, M. Nitta, E.V. Ivanova, R.T. Bronson, and D. Pellman. 2005. Cytokinesis failure generating tetraploids promotes tumorigenesis in p53-null cells. Nature. 437:1043–1047.

Ganem, N.J., Z. Storchova, and D. Pellman. 2007. Tetraploidy, aneuploidy and cancer. Curr Opin Genet Dev. 17:157–162.

Glotzer, M. 2005. The molecular requirements for cytokinesis. Science. 307:1735–1739.

Gluszek-Kustusz, A., B. Craske, T. Legal, T. McHugh, and J.P. Welburn. 2023. Phosphorylation controls spatial and temporal activities of motor-PRC1 complexes to complete mitosis. EMBO J. 42:e113647.

Guizetti, J., L. Schermelleh, J. Mantler, S. Maar, I. Poser, H. Leonhardt, T. Muller-Reichert, and D.W. Gerlich. 2011. Cortical constriction during abscission involves helices of ESCRT-III-dependent filaments. Science. 331:1616–1620.

Halcrow, E.F.J., R. Mazza, A. Diversi, A. Enright, and P.P. D’Avino. 2022. Midbody Proteins Display Distinct Dynamics during Cytokinesis. Cells-Basel. 11.

Hirokawa, N., Y. Noda, Y. Tanaka, and S. Niwa. 2009. Kinesin superfamily motor proteins and intracellular transport. Nat Rev Mol Cell Biol. 10:682–696.

Hosea, R., S. Hillary, S. Naqvi, S. Wu, and V. Kasim. 2024. The two sides of chromosomal instability: drivers and brakes in cancer. Signal Transduct Target Ther. 9:75.

Hu, C.K., M. Coughlin, and T.J. Mitchison. 2012. Midbody assembly and its regulation during cytokinesis. Mol Biol Cell. 23:1024–1034.

Hutterer, A., M. Glotzer, and M. Mishima. 2009. Clustering of centralspindlin is essential for its accumulation to the central spindle and the midbody. Curr Biol. 19:2043–2049.

Iarovaia, O.V., E.P. Minina, E.V. Sheval, D. Onichtchouk, S. Dokudovskaya, S.V. Razin, and Y.S. Vassetzky. 2019. Nucleolus: A Central Hub for Nuclear Functions. Trends Cell Biol. 29:647–659.

Kawashima, T., Y.C. Bao, Y. Nomura, Y. Moon, Y. Tonozuka, Y. Minoshima, T. Hatori, A. Tsuchiya, M. Kiyono, T. Nosaka, H. Nakajima, D.A. Williams, and T. Kitamura. 2006. Rac1 and a GTPase-activating protein, MgcRacGAP, are required for nuclear translocation of STAT transcription factors. J Cell Biol. 175:937–946.

Kosugi, S., M. Hasebe, M. Tomita, and H. Yanagawa. 2009. Systematic identification of cell cycle-dependent yeast nucleocytoplasmic shuttling proteins by prediction of composite motifs. Proc Natl Acad Sci U S A. 106:10171–10176.

Kurasawa, Y., W.C. Earnshaw, Y. Mochizuki, N. Dohmae, and K. Todokoro. 2004. Essential roles of KIF4 and its binding partner PRC1 in organized central spindle midzone formation. EMBO J. 23:3237–3248.

Lawrence, C.J., R.K. Dawe, K.R. Christie, D.W. Cleveland, S.C. Dawson, S.A. Endow, L.S. Goldstein, H.V. Goodson, N. Hirokawa, J. Howard, R.L. Malmberg, J.R. McIntosh, H. Miki, T.J. Mitchison, Y. Okada, A.S. Reddy, W.M. Saxton, M. Schliwa, J.M. Scholey, R.D. Vale, C.E. Walczak, and L. Wordeman. 2004. A standardized kinesin nomenclature. J Cell Biol. 167:19–22.

Lee, Y.M., S. Lee, E. Lee, H. Shin, H. Hahn, W. Choi, and W. Kim. 2001. Human kinesin superfamily member 4 is dominantly localized in the nuclear matrix and is associated with chromosomes during mitosis. Biochem J. 360:549–556.

Lekomtsev, S., K.C. Su, V.E. Pye, K. Blight, S. Sundaramoorthy, T. Takaki, L.M. Collinson, P. Cherepanov, N. Divecha, and M. Petronczki. 2012. Centralspindlin links the mitotic spindle to the plasma membrane during cytokinesis. Nature. 492:276–279.

Lens, S.M.A., and R.H. Medema. 2019. Cytokinesis defects and cancer. Nat Rev Cancer. 19:32–45.

Levesque, A.A., and D.A. Compton. 2001. The chromokinesin Kid is necessary for chromosome arm orientation and oscillation, but not congression, on mitotic spindles. J Cell Biol. 154:1135–1146.

Levine, M.S., and A.J. Holland. 2018. The impact of mitotic errors on cell proliferation and tumorigenesis. Genes Dev. 32:620–638.

Li, T.F., H.J. Zeng, Z. Shan, R.Y. Ye, T.Y. Cheang, Y.J. Zhang, S.H. Lu, Q. Zhang, N. Shao, and Y. Lin. 2020. Overexpression of kinesin superfamily members as prognostic biomarkers of breast cancer. Cancer Cell Int. 20:123.

Liem, K.F., Jr., M. He, P.J. Ocbina, and K.V. Anderson. 2009. Mouse Kif7/Costal2 is a cilia-associated protein that regulates Sonic hedgehog signaling. Proc Natl Acad Sci U S A. 106:13377–13382.

Liu, Y., H. Chen, P. Dong, G. Xie, Y. Zhou, Y. Ma, X. Yuan, J. Yang, L. Han, L. Chen, and L. Shen. 2020. KIF23 activated Wnt/beta-catenin signaling pathway through direct interaction with Amer1 in gastric cancer. Aging (Albany NY). 12:8372–8396.

Lumb, J.H., J.W. Connell, R. Allison, and E. Reid. 2012. The AAA ATPase spastin links microtubule severing to membrane modelling. Biochim Biophys Acta. 1823:192–197.

Mallet, J.D., M.M. Dorr, M.C. Drigeard Desgarnier, N. Bastien, S.P. Gendron, and P.J. Rochette. 2016. Faster DNA Repair of Ultraviolet-Induced Cyclobutane Pyrimidine Dimers and Lower Sensitivity to Apoptosis in Human Corneal Epithelial Cells than in Epidermal Keratinocytes. PLoS One. 11:e0162212.

Martinez-Garay, I., A. Rustom, H.H. Gerdes, and K. Kutsche. 2006. The novel centrosomal associated protein CEP55 is present in the spindle midzone and the midbody. Genomics. 87:243–253.

Maruyama, Y., M. Sugawa, S. Yamaguchi, T. Davies, T. Osaki, T. Kobayashi, M. Yamagishi, S. Takeuchi, M. Mishima, and J. Yajima. 2021. CYK4 relaxes the bias in the off-axis motion by MKLP1 kinesin-6. Commun Biol. 4:180.

Mazumdar, M., J.H. Lee, K. Sengupta, T. Ried, S. Rane, and T. Misteli. 2006. Tumor formation via loss of a molecular motor protein. Curr Biol. 16:1559–1564.

Mazumdar, M., and T. Misteli. 2005. Chromokinesins: multitalented players in mitosis. Trends Cell Biol. 15:349–355.

Mazumdar, M., S. Sundareshan, and T. Misteli. 2004. Human chromokinesin KIF4A functions in chromosome condensation and segregation. J Cell Biol. 166:613–620.

Mazzagatti, A., J.L. Engel, and P. Ly. 2024. Boveri and beyond: Chromothripsis and genomic instability from mitotic errors. Mol Cell. 84:55–69.

Miki, H., M. Setou, K. Kaneshiro, and N. Hirokawa. 2001. All kinesin superfamily protein, KIF, genes in mouse and human. Proc Natl Acad Sci U S A. 98:7004–7011.

Morita, E., V. Sandrin, H.Y. Chung, S.G. Morham, S.P. Gygi, C.K. Rodesch, and W.I. Sundquist. 2007. Human ESCRT and ALIX proteins interact with proteins of the midbody and function in cytokinesis. EMBO J. 26:4215–4227.

Pan, H., R. Guan, R. Zhao, G. Ou, and Z. Chen. 2021. Mechanistic insights into central spindle assembly mediated by the centralspindlin complex. Proc Natl Acad Sci U S A. 118.

Park, H., M. Choi, Y. Zhang, H.O. Cheung, S. Makino, Y. Yoshikawa, H. Qi, Z. Liu, G. Lan, G. Fu, Q. Wang, S.S. Guo, P. Liu, Z. Liu, S.C. Ti, W.J. Wang, X.D. Li, T. Ni, C.C. Hui, and M. He. 2025. The kinesin-4 protein KIF27 forms a cytoskeletal scaffold at the transition zone to promote motile cilia structural integrity. Proc Natl Acad Sci U S A. 122:e2515392122.

Petsalaki, E., and G. Zachos. 2016. Clks 1, 2 and 4 prevent chromatin breakage by regulating the Aurora B-dependent abscission checkpoint. Nat Commun. 7:11451.

Pust, S., A. Brech, C.S. Wegner, H. Stenmark, and K. Haglund. 2023. Vesicle-mediated transport of ALIX and ESCRT-III to the intercellular bridge during cytokinesis. Cell Mol Life Sci. 80:235.

Radulovic, M., K.O. Schink, E.M. Wenzel, V. Nahse, A. Bongiovanni, F. Lafont, and H. Stenmark. 2018. ESCRT-mediated lysosome repair precedes lysophagy and promotes cell survival. EMBO J. 37.

Ramkumar, N., J.V. Patel, J. Anstatt, and B. Baum. 2021. Aurora B-dependent polarization of the cortical actomyosin network during mitotic exit. EMBO Rep:e52387.

Scheffler, J.M., N. Schiefermeier, and L.A. Huber. 2014. Mild fixation and permeabilization protocol for preserving structures of endosomes, focal adhesions, and actin filaments during immunofluorescence analysis. Methods Enzymol. 535:93–102.

Schindelin, J., I. Arganda-Carreras, E. Frise, V. Kaynig, M. Longair, T. Pietzsch, S. Preibisch, C. Rueden, S. Saalfeld, B. Schmid, J.Y. Tinevez, D.J. White, V. Hartenstein, K. Eliceiri, P. Tomancak, and A. Cardona. 2012. Fiji: an open-source platform for biological-image analysis. Nature methods. 9:676–682.

Sisson, J.C., K.S. Ho, K. Suyama, and M.P. Scott. 1997. Costal2, a novel kinesin-related protein in the Hedgehog signaling pathway. Cell. 90:235–245.

Smith, E.R., C.D. Capo-Chichi, and X.X. Xu. 2018. Defective Nuclear Lamina in Aneuploidy and Carcinogenesis. Front Oncol. 8:529.

Steigemann, P., C. Wurzenberger, M.H. Schmitz, M. Held, J. Guizetti, S. Maar, and D.W. Gerlich. 2009. Aurora B-mediated abscission checkpoint protects against tetraploidization. Cell. 136:473–484.

Stirling, D.R., M.J. Swain-Bowden, A.M. Lucas, A.E. Carpenter, B.A. Cimini, and A. Goodman. 2021. CellProfiler 4: improvements in speed, utility and usability. BMC Bioinformatics. 22:433.

Takahashi, M., T. Wakai, and T. Hirota. 2016. Condensin I-mediated mitotic chromosome assembly requires association with chromokinesin KIF4A. Genes Dev. 30:1931–1936.

Tanaka, K., and T. Hirota. 2009. Chromosome segregation machinery and cancer. Cancer Sci. 100:1158–1165.

Tang, F., M.H. Pan, Y. Lu, X. Wan, Y. Zhang, and S.C. Sun. 2018. Involvement of Kif4a in Spindle Formation and Chromosome Segregation in Mouse Oocytes. Aging Dis. 9:623–633.

Tokai-Nishizumi, N., M. Ohsugi, E. Suzuki, and T. Yamamoto. 2005. The chromokinesin Kid is required for maintenance of proper metaphase spindle size. Mol Biol Cell. 16:5455–5463.

Vanneste, D., V. Ferreira, and I. Vernos. 2011. Chromokinesins: localization-dependent functions and regulation during cell division. Biochem Soc Trans. 39:1154–1160.

Vasudevan, A., K.M. Schukken, E.L. Sausville, V. Girish, O.A. Adebambo, and J.M. Sheltzer. 2021. Aneuploidy as a promoter and suppressor of malignant growth. Nat Rev Cancer. 21:89–103.

Vicidomini, G., P. Bianchini, and A. Diaspro. 2018. STED super-resolved microscopy. Nature methods. 15:173–182.

Vietri, M., M. Radulovic, and H. Stenmark. 2020. The many functions of ESCRTs. Nat Rev Mol Cell Biol. 21:25–42.

Wandke, C., M. Barisic, R. Sigl, V. Rauch, F. Wolf, A.C. Amaro, C.H. Tan, A.J. Pereira, U. Kutay, H. Maiato, P. Meraldi, and S. Geley. 2012. Human chromokinesins promote chromosome congression and spindle microtubule dynamics during mitosis. J Cell Biol. 198:847–863.

Wang, S.Z., and R. Adler. 1995. Chromokinesin: a DNA-binding, kinesin-like nuclear protein. J Cell Biol. 128:761–768.

White, E.A., and M. Glotzer. 2012. Centralspindlin: at the heart of cytokinesis. Cytoskeleton (Hoboken). 69:882–892.

Wollman, A.J., C. Sanchez-Cano, H.M. Carstairs, R.A. Cross, and A.J. Turberfield. 2014. Transport and self-organization across different length scales powered by motor proteins and programmed by DNA. Nat Nanotechnol. 9:44–47.

Yildiz, A. 2025. Mechanism and regulation of kinesin motors. Nat Rev Mol Cell Biol. 26:86–103.

Yue, Y., T.L. Blasius, S. Zhang, S. Jariwala, B. Walker, B.J. Grant, J.C. Cochran, and K.J. Verhey. 2018. Altered chemomechanical coupling causes impaired motility of the kinesin-4 motors KIF27 and KIF7. J Cell Biol. 217:1319–1334.

Zhao, W.M., A. Seki, and G. Fang. 2006. Cep55, a microtubule-bundling protein, associates with centralspindlin to control the midbody integrity and cell abscission during cytokinesis. Mol Biol Cell. 17:3881–3896.

Zhong, A., F.Q. Tan, and W.X. Yang. 2016. Chromokinesin: Kinesin superfamily regulating cell division through chromosome and spindle. Gene. 589:43–48.

