## Supplemental Figures and Tables for "The kinesin-4 family member KIF27 regulates mitotic progression, cytokinesis and genome stability"

### suppl. Fig. 1

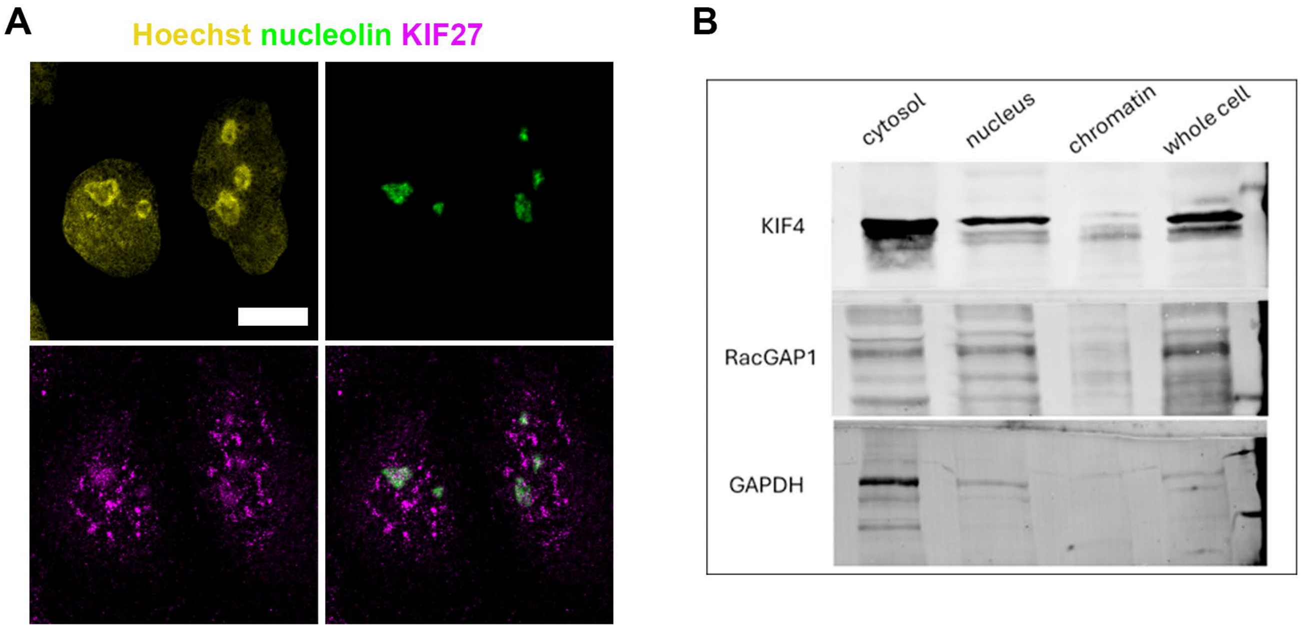


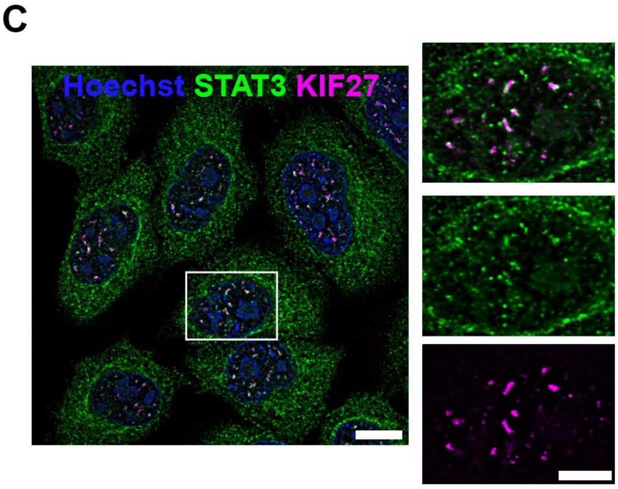


**D**

MEEIPVKVAVRIRPLLCKEALHNHQVCVRVIPNSQQVIIGRDRVFTFDFVFGKNSTQDEV
YNTCIKPLVLSLIEGYNATVFAYGQTGSGKTYTIGGGHIASVVEGQKGIIPRAIQEIFQS
ISEHPSIDFNVKVSYIEVYKEDLRDLLELETSMKDLHIREDEKGNTVIVGAKECHVESAG
EVMSLLEMGNAARHTGTTQMNEHSSRSHAIFTISICQVHKNMEAAEDGSWYSPRHIVSKF
HFVDLAGSERVTKTGNTGERFKESIQINSGLLALGNVISALGDPRRKSSHIPYRDAKITR
LLKDSLGGSAKTVMITCVSPSSSNFDESLNSLKYANRARNIRNKPTVNFSPESDRIDEME
FEIKLLREALQSQQAGVSQTTQINREGSPDTNRIHSLEEQVAQLQGECLGYQCCVEEAFT
FLVDLKDTVRLNEKQQHKLQEWFNMIQEVRKAVLTSFRGIGGTASLEEGPQHVTVLQLKR
ELKKCQCVLAADEVVFNQKELEVKELKNQVQMMVQENKGHAVSLKEAQKVNRLQNEKIIE
QQLLVDQLSEELTKLNLSVTSSAKENCGDGPDARIPERRPYTVPFDTHLGHYIYIPSRQD
SRKVHTSPPMYSLDRIFAGFRTRSQMLLGHIEEQDKVLHCQFSDNSDDEESEGQEKSGTR
CRSRSWIQKPDSVCSLVELSDTQDETQKSDLENEDLKIDCLQESQELNLQKLKNSERILT
EAKQKMRELTINIKMKEDLIKELIKTGNDAKSVSKQYSLKVTKLEHDAEQAKVELIETQK
QLQELENKDLSDVAMKVKLQKEFRKKMDAAKLRVQVLQKKQQDSKKLASLSIQNEKRANE
LEQSVDHMKYQKIQLQRKLREENEKRKQLDAVIKRDQQKIKEIQLKTGQEEGLKPKAEDL
DACNLKRRKGSFGSIDHLQKLDEQKKWLDEEVEKVLNQRQELEELEADLKKREAIVSKKE
ALLQEKSHLENKKLRSSQALNTDSLKISTRLNLLEQELSEKNVQLQTSTAEEKTKISEQV
EVLQKEKDQLQKRRHNVDEKLKNGRVLSPEEEHVLFQLEEGIEALEAAIEYRNESIQNRQ
KSLRASFHNLSRGEANVLEKLACLSPVEIRTILFRYFNKVVNLREAERKQQLYNEEMKMK
VLERDNMVRELESALDHLKLQCDRRLTLQQKEHEQKMQLLLHHFKEQDGEGIMETFKTYE
DKIQQLEKDLYFYKKTSRDHKKKLKELVGEAIRRQLAPSEYQEAGDGVLKPEGGGMLSEE
LKWASRPESMKLSGREREMDSSASSLRTQPNPQKLWEDIPELPPIHSSLAPPSGHMLGNE
NKTETDDNQFTKSHSRLSSQIQVVGNVGRLHGVTPVKLCRKELRQISALELSLRRSSLGV
GIGSMAADSIEVSRKPRDLKT

suppl. Fig.2

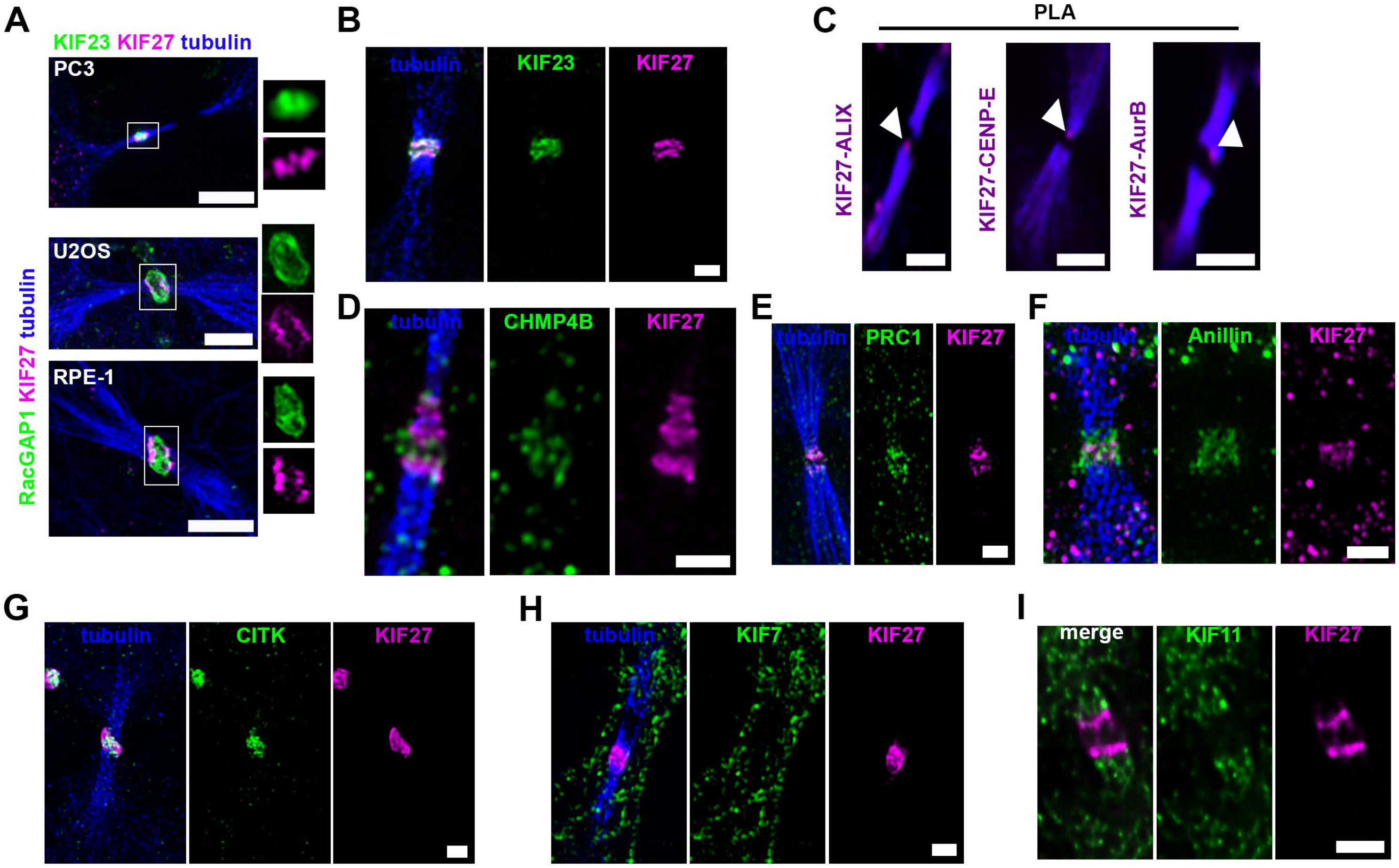
A
KIF23 KIF27 tubulin
PC3
U2OS
RPE-1
RacGAP1 KIF27 tubulin
B
tubulin
KIF23
KIF27
C
PLA
KIF27-ALIX
KIF27-CENP-E
KIF27-AurB
D
tubulin
CHMP4B
KIF27
E
tubulin
PRC1
KIF27
F
tubulin
Anillin
KIF27
G
tubulin
CITK
KIF27
H
tubulin
KIF7
KIF27
I
merge
KIF11
KIF27

**suppl. Fig. 3**

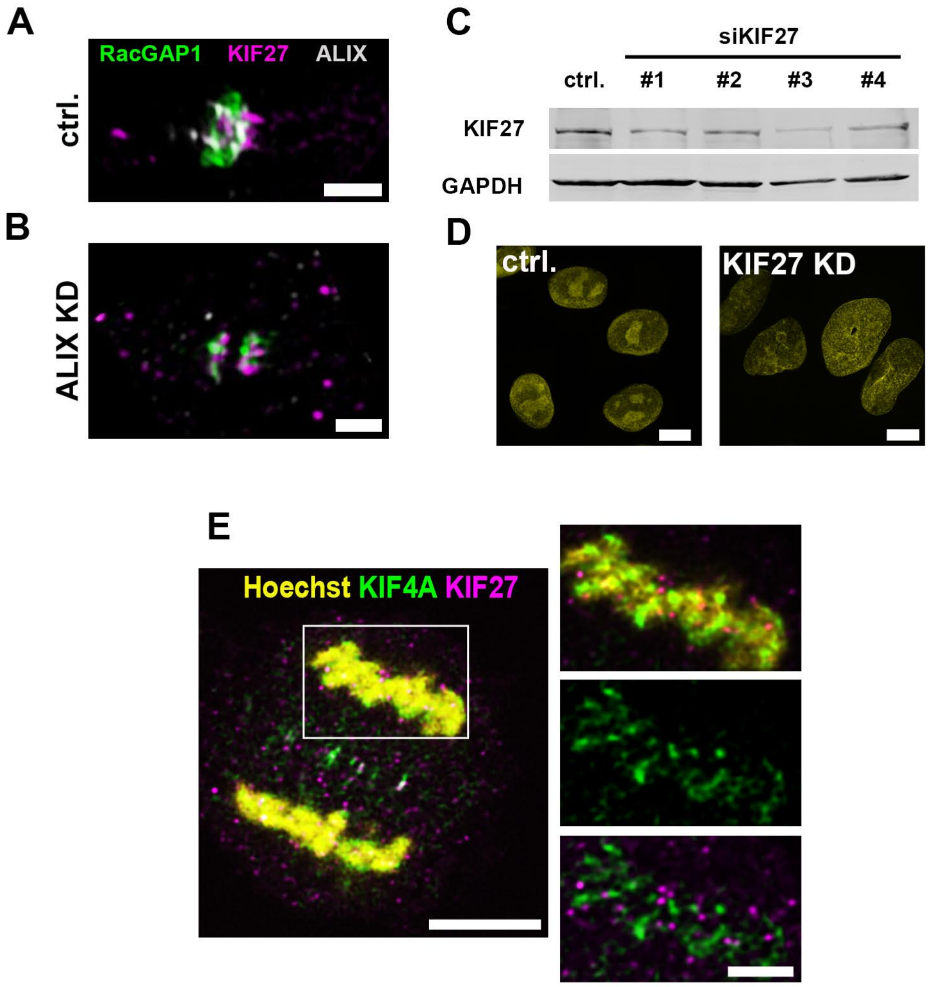
A
RacGAP1 KIF27 ALIX
ctrl.
B
ALIX KD
C
siKIF27
ctrl. #1 #2 #3 #4
KIF27
GAPDH
D
ctrl.
KIF27 KD
E
Hoechst KIF4A KIF27

**suppl. Fig. 4**

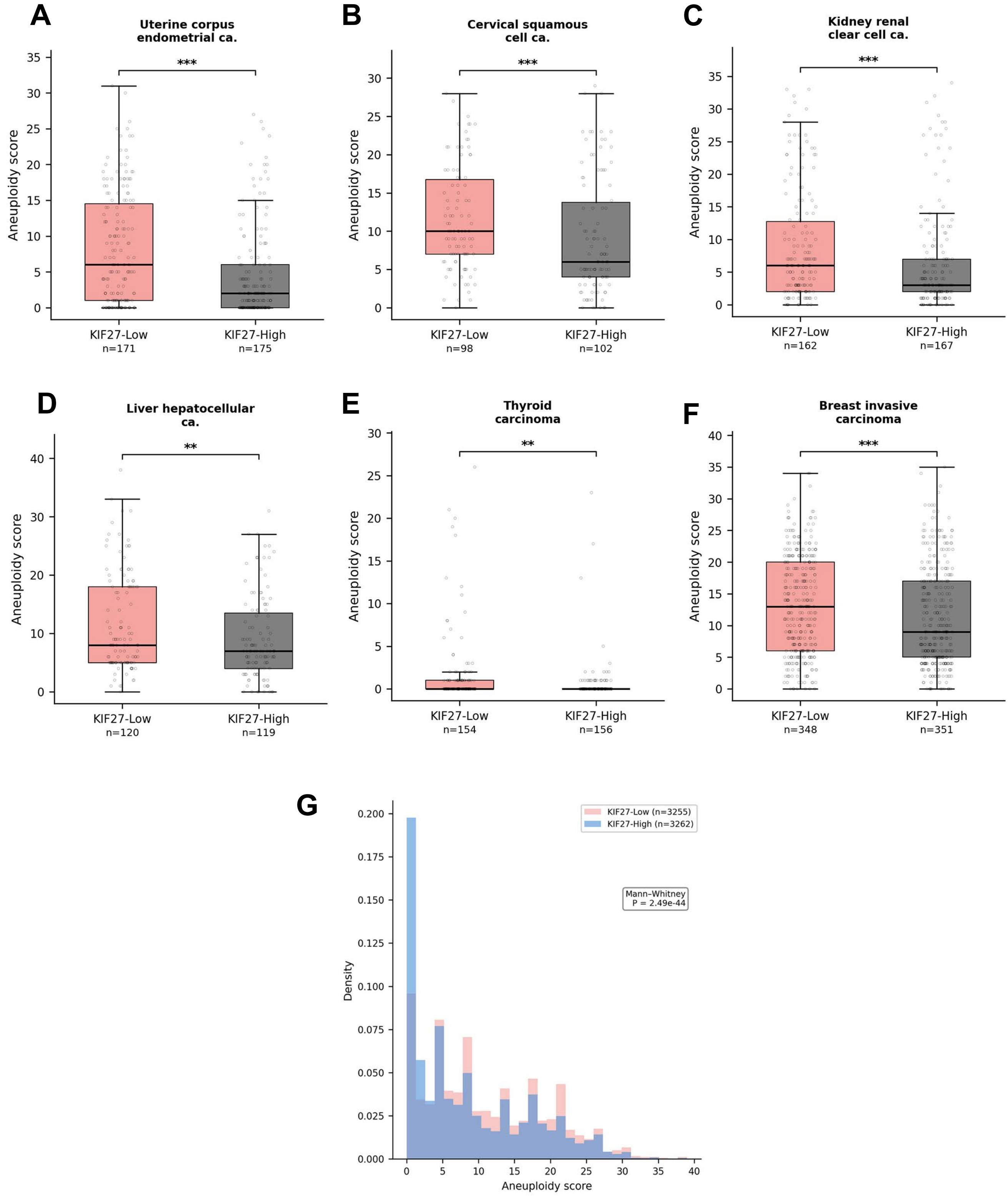

A
Uterine corpus endometrial ca.
***
Aneuploidy score
KIF27-Low n=171
KIF27-High n=175
B
Cervical squamous cell ca.
***
Aneuploidy score
KIF27-Low n=98
KIF27-High n=102
C
Kidney renal clear cell ca.
***
Aneuploidy score
KIF27-Low n=162
KIF27-High n=167
D
Liver hepatocellular ca.
**
Aneuploidy score
KIF27-Low n=120
KIF27-High n=119
E
Thyroid carcinoma
**
Aneuploidy score
KIF27-Low n=154
KIF27-High n=156
F
Breast invasive carcinoma
***
Aneuploidy score
KIF27-Low n=348
KIF27-High n=351
G
KIF27-Low (n=3255)
KIF27-High (n=3262)
Mann-Whitney
P = 2.49e-44
Density
Aneuploidy score

suppl. Fig. 5

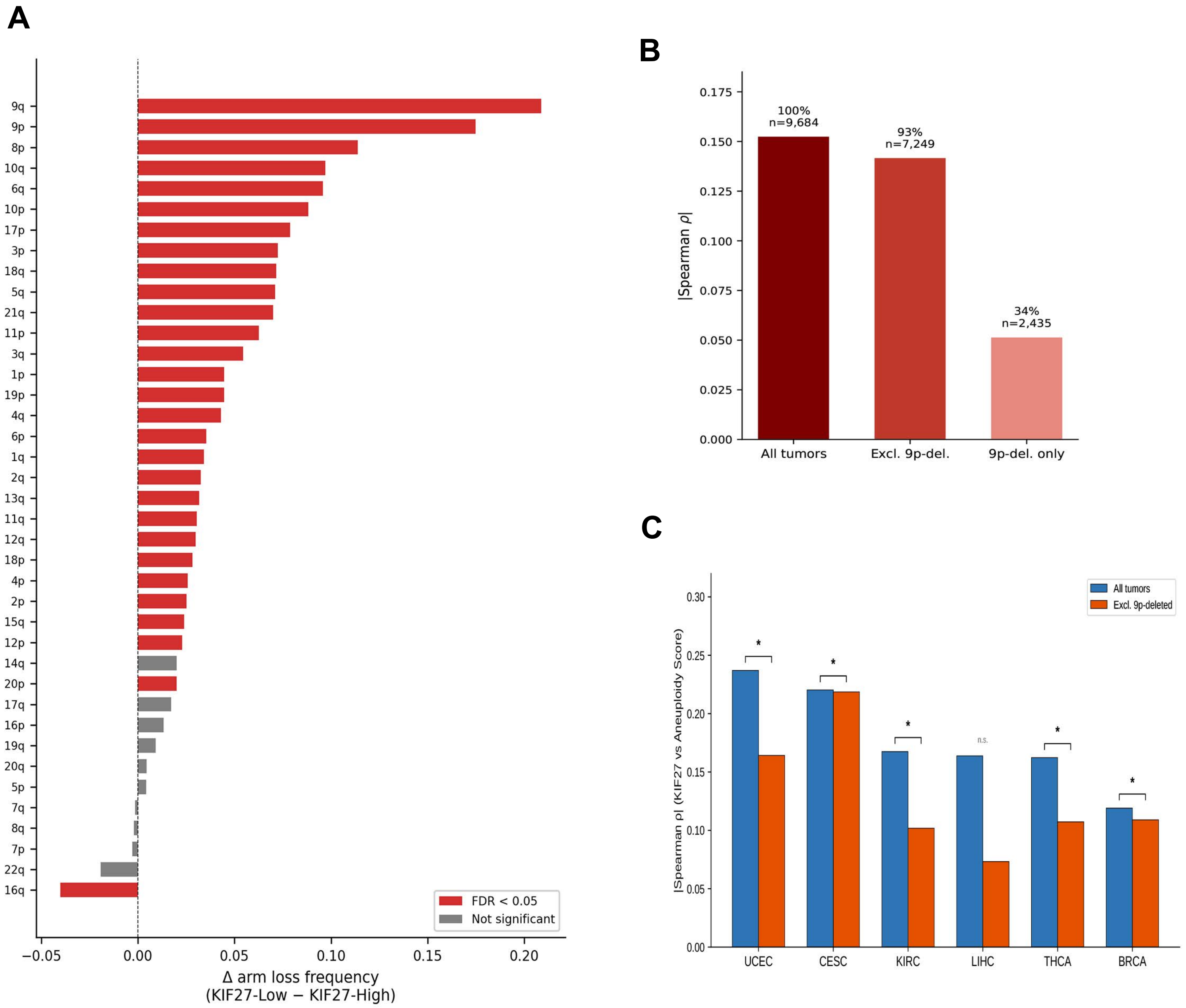
A
9q
9p
8p
10q
6q
10p
17p
3p
18q
5q
21q
11p
3q
1p
19p
4q
6p
1q
2q
13q
11q
12q
18p
4p
2p
15q
12p
14q
20p
17q
16p
19q
20q
5p
7q
8q
7p
22q
16q
FDR < 0.05
Not significant
−0.05
0.00
0.05
0.10
0.15
0.20
Δ arm loss frequency
(KIF27-Low − KIF27-High)
B
|Spearman ρ|
0.175
0.150
0.125
0.100
0.075
0.050
0.025
0.000
100%
n=9,684
93%
n=7,249
34%
n=2,435
All tumors
Excl. 9p-del.
9p-del. only
C
|Spearman ρ| (KIF27 vs Aneuploidy Score)
0.30
0.25
0.20
0.15
0.10
0.05
0.00
All tumors
Excl. 9p-deleted
n.s.
UCEC
CESC
KIRC
LIHC
THCA
BRCA

#### Supplementary Table S1

| Cancer type | n | ρ | P-value | FDR q | Sig. |
|---|---|---|---|---|---|
| Uveal melanoma (UVM) | 80 | -0.273 | 0.014 | 0.068 | |
| **Uterine corpus endometrial ca. (UCEC)** | **514** | **-0.237** | **5.4 $\times 10^{-08}$** | **1.8 $\times 10^{-06}$** | ✓ |
| **Cervical squamous cell ca. (CESC)** | **291** | **-0.220** | **1.5 $\times 10^{-04}$** | **0.002** | ✓ |
| **Kidney renal clear cell ca. (KIRC)** | **481** | **-0.168** | **2.2 $\times 10^{-04}$** | **0.002** | ✓ |
| **Liver hepatocellular ca. (LIHC)** | **355** | **-0.164** | **0.002** | **0.011** | ✓ |
| **Thyroid carcinoma (THCA)** | **462** | **-0.162** | **4.6 $\times 10^{-04}$** | **0.003** | ✓ |
| Rectum adenocarcinoma (READ) | 154 | -0.132 | 0.103 | 0.263 | |
| **Breast invasive ca. (BRCA)** | **1039** | **-0.119** | **1.2 $\times 10^{-04}$** | **0.002** | ✓ |
| Stomach adenocarcinoma (STAD) | 401 | -0.114 | 0.022 | 0.091 | |
| Uterine carcinosarcoma (UCS) | 56 | -0.108 | 0.427 | 0.611 | |
| Kidney chromophobe (KICH) | 65 | -0.095 | 0.451 | 0.611 | |
| Acute myeloid leukemia (LAML) | 109 | -0.085 | 0.379 | 0.569 | |
| Kidney renal papillary cell ca. (KIRP) | 280 | -0.079 | 0.186 | 0.410 | |
| Lung adenocarcinoma (LUAD) | 497 | -0.074 | 0.099 | 0.263 | |
| Ovarian serous cystadenocarcinoma (OV) | 289 | -0.066 | 0.265 | 0.453 | |
| Thymoma (THYM) | 103 | -0.054 | 0.589 | 0.748 | |
| Colon adenocarcinoma (COAD) | 430 | -0.053 | 0.275 | 0.453 | |
| Lung squamous cell ca. (LUSC) | 479 | -0.034 | 0.463 | 0.611 | |
| Head & neck squamous cell ca. (HNSC) | 502 | -0.009 | 0.844 | 0.871 | |
| Pancreatic adenocarcinoma (PAAD) | 158 | 0.001 | 0.993 | 0.993 | |
| Testicular germ cell tumors (TGCT) | 149 | 0.018 | 0.824 | 0.871 | |
| Mesothelioma (MESO) | 81 | 0.024 | 0.835 | 0.871 | |
| Glioblastoma multiforme (GBM) | 152 | 0.030 | 0.716 | 0.844 | |
| Adrenocortical ca. (ACC) | 76 | 0.030 | 0.798 | 0.871 | |
| Esophageal ca. (ESCA) | 161 | 0.032 | 0.688 | 0.841 | |
| Brain lower grade glioma (LGG) | 508 | 0.053 | 0.237 | 0.453 | |
| Prostate adenocarcinoma (PRAD) | 471 | 0.068 | 0.142 | 0.335 | |
| Skin cutaneous melanoma (SKCM) | 458 | 0.085 | 0.070 | 0.210 | |
| Pheochromocytoma & paraganglioma (PCPG) | 160 | 0.097 | 0.221 | 0.453 | |
| Bladder urothelial ca. (BLCA) | 398 | 0.104 | 0.038 | 0.140 | |
| Sarcoma (SARC) | 242 | 0.125 | 0.052 | 0.172 | |
| Cholangiocarcinoma (CHOL) | 36 | 0.153 | 0.374 | 0.569 | |
| Diffuse large B-cell lymphoma (DLBC) | 47 | 0.172 | 0.247 | 0.453 | |

#### Supplementary Table S2

| Cancer type | n | Events | KIF27 → OS (KMplot HR) | KIF27 → OS KMplot p | AS → OS (TCGA HR) | AS → OS 95% CI | AS → OS Cox p | Sig. |
|---|---|---|---|---|---|---|---|---|
| **Esoph. SCC** | 84 | 30 | HR = 0.33 (favorable) | 0.0140 | 1.002 | (0.953–1.052) | 0.9500 | ns |
| **HNSC** | 511 | 213 | HR = 0.66 (favorable) | 0.0120 | 1.015 | (0.997–1.033) | 0.1109 | ns |
| **KIRC** | 481 | 164 | HR = 0.65 (favorable) | 0.0047 | 1.012 | (0.995–1.030) | 0.1690 | ns |
| **LUAD** | 491 | 172 | HR = 0.72 (favorable) | 0.0450 | 1.008 | (0.989–1.028) | 0.4084 | ns |
| **PCPG** | 164 | 7 | HR = 9.05 (unfavorable) | 0.0180 | 0.986 | (0.841–1.155) | 0.8596 | ns |
| **Rectum** | 148 | 22 | HR = 0.17 (favorable) | 0.0072 | 0.991 | (0.938–1.047) | 0.7519 | ns |
| **Sarcoma** | **248** | **93** | **HR = 1.72 (unfavorable)** | **0.0084** | **1.029** | **(1.005–1.054)** | **0.0182** | * |
